# Rewiring Cancer Drivers to Activate Apoptosis

**DOI:** 10.1101/2022.12.04.517548

**Authors:** Sai Gourisankar, Andrey Krokhotin, Wenzhi Ji, Xiaofan Liu, Chiung-ying Chang, Samuel H. Kim, Zhengnian Li, Wendy Wenderski, Juste M. Simanauskaite, Tinghu Zhang, Nathanael S. Gray, Gerald R. Crabtree

## Abstract

Genes that drive the proliferation, survival, invasion and metastasis of malignant cells have been identified for many human cancers^1–6^. Independent studies have identified cell death pathways that eliminate cells for the good of the organism^7–10^. The coexistence of the cell death pathways with the driver mutations suggest that the cancer driver could be rewired to activate cell death. We have invented a new class of molecules: TCIPs (Transcriptional/Epigenetic Chemical Inducers of Proximity) that recruit the endogenous cancer driver, or a downstream transcription factor, to the promoters of cell death genes thereby activating their expression. To develop this concept, we have focused on diffuse large B cell lymphoma (DLBCL), in which BCL6 is amplified or mutated^11^. BCL6 binds to the promoters of cell death genes and epigenetically suppresses their expression^12^. We produced the first TCIPs by chemically linking BCL6 inhibitors to small molecules that bind transcriptional activators. Several of these molecules robustly kill DLBCL at single-digit nanomolar concentrations, including chemotherapy-resistant, TP53-mutant lines. The dominant gain-of-function approach provided by TCIPs captures the combinatorial specificity inherit to transcription and can thereby accesses new therapeutic space. TCIPs are relatively non-toxic to normal cells and mice, apparently reflecting their need for coincident expression of both target proteins for effective killing. The general TCIP concept has applications in elimination of senescent cells, enhancing expression of therapeutic genes, treatment of diseases produced by haploinsufficiency, and activation of immunogens for immunotherapy.

## TEXT

Chemically induced proximity (CIP) underlies many forms of biologic regulation. These include post translational modifications such as phosphorylation and methylation as well as allosteric processes that generate platforms or scaffolds that facilitate protein interactions. The biologic roles of chemically induced proximity have been probed with genetically tagged proteins that allow resolution of the roles of proximity and conformational modifications^13–17^. More recently, small molecules that use CIP to target proteins to the proteasome, PROTACS^18^, or to induce stabilization^19^ have been developed. These studies have revealed that chemically induced proximity underlies, or at least can reproduce many steps in signal transduction and transcription^20–23^. In the nucleus induced proximity is essential for several steps in the activation of transcription^22^ as well as epigenetic repression or activation^24–26^.

An informative example of the role of CIP is the activation of death receptors such as the FAS receptor (FASR or CD95) which eliminate autoreactive cells in the immune system. Induced proximity of their death domains or of the initiator caspases^27,28^ rapidly and robustly initiates apoptosis^14,29–31^. The observation that an event as carefully regulated as programmed cell death can be activated by CIP suggests that members of these signal transduction and transcriptional pathways might be linked or rewired to cause cancer cells to activate processes leading to apoptosis.

### TCIP1 selectively kills DLBCL cells at low nanomolar concentrations

To rewire transcriptional circuits within a genetically unmodified cell or organism we have developed molecules that allow the recruitment of transcriptional or epigenetic regulators to the regulatory regions of target therapeutic genes. The general features of the concept and design of a TCIP is illustrated in Fig. 1a and involves synthesis of small molecules that bind a specific transcription factor on one side, and on the other side, a transcription factor that binds to a target therapeutic gene.

**Figure 1.**
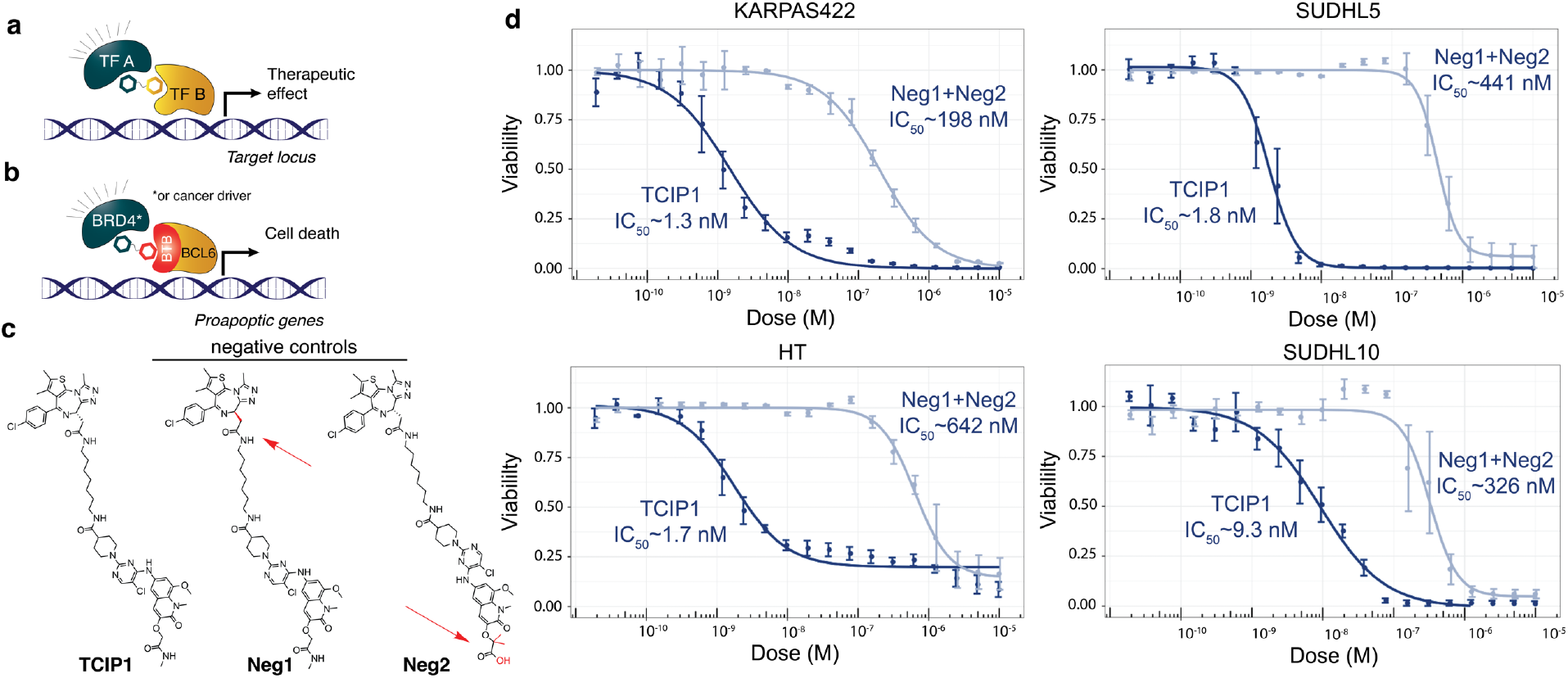
Transcriptional Chemical Inducers of Proximity (TCIPs) to rewire transcriptional circuits. **a.** An endogenous target gene is activated or repressed using a bifunctional molecule binding one endogenous transcription factor or epigenetic regulator on one side chemically linked to a moiety that binds a second transcription factor binding the regulatory region of a target gene which might produce a therapeutic target. **b.** A specific TCIP that recruits a transcriptional activator (BRD4) or cancer driver to the BCL6 repressor on cell death genes, thereby derepressing transcription and inducing transcription driven by BCL6. **c.** Chemical structures of the most potent BCL6/BRD4 TCIP, TCIP1, and negative controls Neg1 (BRD4 non-binding) and Neg2 (BCL6 non-binding) **d.** TCIP1 effect on cell viability of the CHOP-resistant, TP53 mutant diffuse large B-cell lymphoma cell (DLBCL) line KARPAS422 as well as 3 other DLBCL cell lines with high levels of BCL6. n=4 biological repeats, mean±s.d.

To design the first TCIPs, we targeted DLBCL and made use of small molecules that bind to the BTB domain of BCL6 and prevent the binding of NCOR, BCOR and SMRT, which epigenetically suppress BCL6 targets including pro-apoptotic, cell cycle arrest, and DNA-damage response genes^32^, such as *TP53*^12^, by binding HDACs^33^ and Polycomb Repressive Complexes (PRC)^34^ (Fig 1b). Although these small molecules derepress BCL6 targets, they and related BCL6 degraders have shown limited eficacy^35,36^. To provide additional transcriptional activation of proapoptotic genes, over simple derepression, we linked the inhibitor BI3812^36^ covalently to JQ1^37^, which binds comparably to both bromodomains of BRD4, a bromodomain protein involved in transcriptional activation, which also contributes a driving function to several tumors by facilitating MYC activation^38^. We synthesized a series of molecules with linkers of different lengths and flexibility including TCIP1 (Fig 1c). These molecules were tested for their effect on viability of the CHOP-resistant DLBCL cell line, KARPAS422. This line has biallelic inactivation of TP53 and was chosen for its high level of expression of BCL6, and the fact that it was derived from a tumor that had become resistant to CHOP therapy and has multiple drivers^39,40^. TCIP1 rapidly and robustly killed KARPAS422 with an EC50 of 1.3 nM, 72 hours after addition of drug. Killing was essentially complete at 72 hours (Fig 1d). Observation of the cultures for 4 weeks identified no clones that escaped killing.

Three other DLBCL lines with high levels of BCL6 (Fig. 1d) were also rapidly and robustly killed by TCIP1. Adding JQ1 and BI3812 separately or together showed 100-fold to 1000-fold less effective cell killing (Ext. Data Fig 1a.), excluding the possibility that TCIP1 acts by simply delivering two inhibitors into the cell. We also synthesized negative chemical controls, Neg1 and Neg2, having the same linker structure as TCIP1 but having modifications known to mitigate binding to BRD4 or BCL6, respectively^36,37^. Neg1 and Neg2 had greater than 100-fold less effect on cell viability compared to TCIP1, even in combination (Fig. 1d), suggesting that binding both proteins in proximity is required for effective killing. We noted that neither BI3812 or JQ1 produced complete killing leaving a resistant population of cells as has been previously reported^41^.We tested a panel of 14 lymphoma and other cancer cell lines expressing different levels of BCL6. As expected, killing as measured by EC50 was correlated with BCL6 levels and BCL6 mRNA levels were predictive of sensitivity to TCIP1 (Ext. Data Fig 1a). Note that only 2/14 (RAJI and HT) of the DLBCL cell lines profiled are unambiguously sensitive to BCL6 CRISPR knockout^42^. Some tumor cell lines such as Jurkat, with no detectable level of BCL6 (Ext. Data Fig 1b), showed little or no response to sub-micromolar concentrations of the drugs. The level of expression of BCOR, NCOR and SMRT, which engage PRC and HDACs to produce epigenetic suppression of target genes^33,34^ varied among the cell lines and could contribute to the variation in sensitivity^43^. We confirmed that protein levels of p53, BCL6, BRD4, and BCL2 in our cell lines are consistent with previous data on their gene expression (Ext. Data Fig. 1b). Among the cell lines tested there was no evidence that killing required TP53, nor was there evidence of repression of killing by endogenous BCL2 levels (Ext. Data Fig. 1b). We noted that TCIP1 was more than 1000-fold as potent in killing DLBCL cells compared to degradation of BRD4 by dBET1^44^ or degradation of BCL6 by BI3802^45^ (Ext. Data Fig. 1c). In an unbiased screen of the effect of TCIP1 on the viability of 906 cancer cell lines (PRISM^46^), almost all sensitive cancer cells originated from hematopoetic and lymphoid tissues, and potency was correlated with BCL6 expression (Ext. Data Fig. 1d).

### Lymphoma cell killing requires formation of a ternary complex

To determine if formation of a ternary complex between BRD4, BCL6, and TCIP1 in cells was essential for effective killing, we carried out chemical rescue experiments in which we titrated increasing concentrations of either JQ1 or BI3812 to multiple DLBCL cell lines against constant concentrations of TCIP1 that kill 50 to 95% of cells within 72 hours. Increasing concentrations of either JQ1 or BI3812 prevented death by TCIP1, indicating that both the BRD4 binding side and the BCL6 binding side of TCIP1 are essential for effective killing (Fig. 2a,b). Examination of other DLBCL lines indicated that rescue was most prominent in highly sensitive cell lines (Ext. Data Fig. 2).

**Figure 2.**
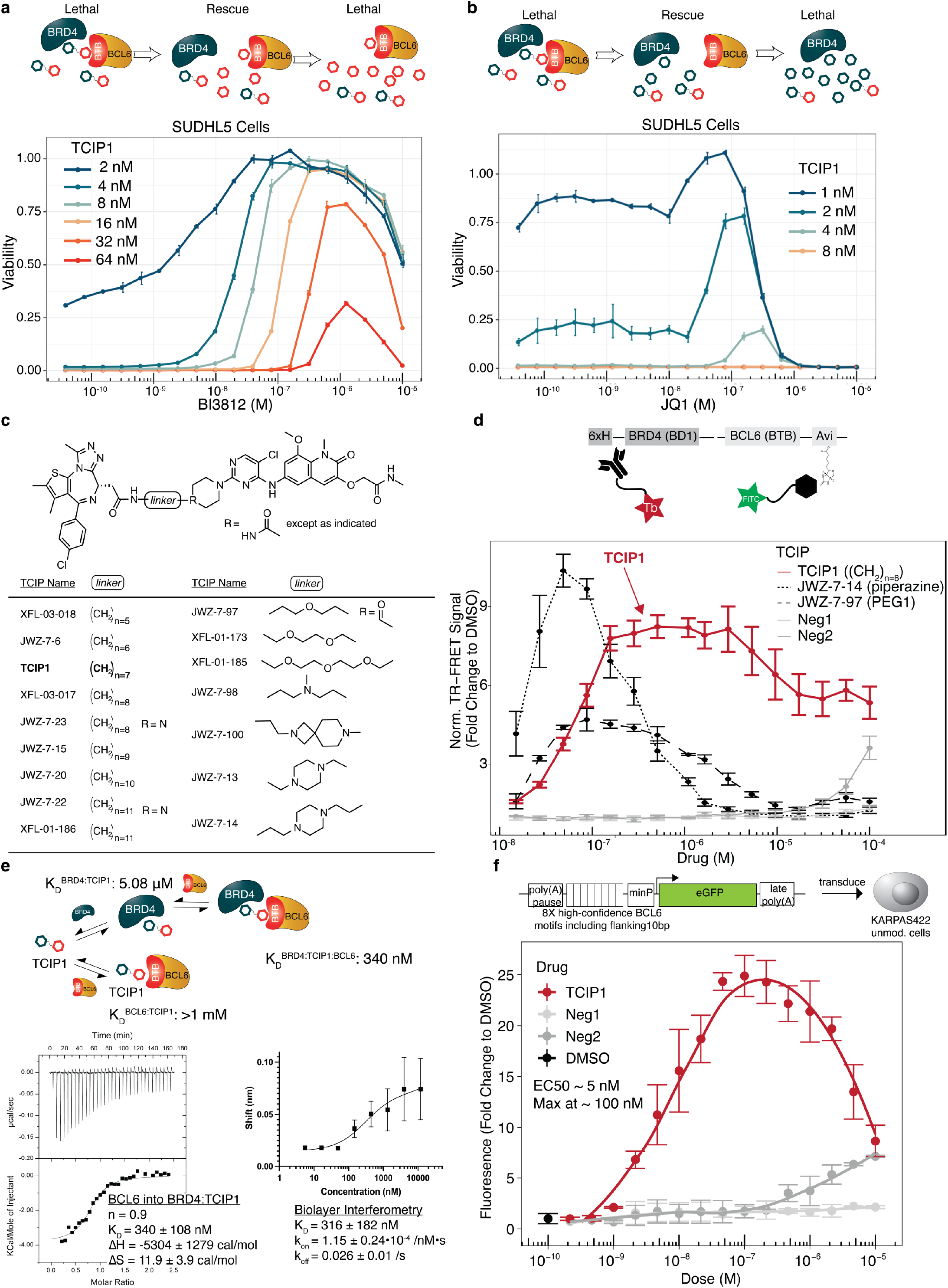
TCIP function requires ternary complex formation. **a.** Competitive titration of BI3812 against TCIP1. TCIP1 was added at concentrations from 2-64 nM that killed 90% of SUDHL5 DLBCL cells at the same time as addition of the indicated concentrations of BI3812. n=3 biological repeats, mean±s.d.. **b.** Competitive titration of JQ1 against TCIP1. n=3 biological repeats, mean±s.d. **c.** Multiple BRD4-BCL6 TCIPs synthesized with different linkers to test structure-activity relationship. **d.** TR-FRET assay to measure molecule-dependent ternary complex formation between BRD4^BD1^ and BCL6^BTB^. Plotted are a representative set of TCIPs that were the most potent (in cell viability assays) within each category of linker structure. TCIP1 had the highest potency of all designed molecules. n=2 independent repeats with different batches of protein with n=3 technical repeats each, mean±s.e. **e.** Analysis of cooperative binding induced by TCIP1 and BRD4^BD1^ and BCL6^BTB^ domains. Representative ternary complex K_D_ measurement by isothermal calorimetry (ITC) shown, for binary measurements see Extended Data Figure 2d. n=3 independent repeats, mean±s.d. ITC parameters shown with 20:1 BRD4^BD1^:TCIP1 in the cell and titration of BCL6^BTB^. For biolayer interferometry (BLI) measurements were with 50μM excess BRD4^BD1^ in the well, nanomolar titrations of TCIP1, and biotinylated BCL6^BTB^ on the tip, n=2 independent repeats, mean±s.d. **f.** Design and activation of a BCL6 reporter with TCIP1 in KARPAS422 cells at 8hrs drug addition. n=4 biological repeats, mean±s.d.

To gain evidence of a direct induced proximity interaction between TCIP1, BRD4, and BCL6, we synthesized several related TCIPs employing different linkers (Fig. 2c) and developed a TR-FRET assay based on the proximity of the BTB domain of BCL6 labeled with FITC to bromodomain 1 of BRD4 detected with an anti-histidine-tag, terbium-conjugated antibody (Fig. 2d). Formation of a ternary complex *in vitro*, related to TR-FRET peak height^47^, was detected using multiple TCIP molecules shown in Fig. 1c. More flexible or longer linkers reduced ternary complex formation and increased the concentration required for the signal to peak (Ext. Data Fig. 2c). TCIP1 was the most potent in cell killing, despite not having the highest affinity ternary complex formation by TR-FRET of the compounds made and tested (Fig. 2d,e). The calculated affinity of the ternary complex maximum (500nM for TCIP1) was less than the EC50 for cell death in the most sensitive DLBCL cell lines. This disparity may reflect the fact that we have used small fragments of the two proteins for *in vitro* FRET detection and hence there would be no contribution of any region of the two proteins outside of the fragments we have used for ternary complex formation, while in cells, interactions between the entire proteins are feasible, such as the second bromodomain of BRD4. Of note is the fact that the signal is a skewed “hook” and does not return to zero at high levels of TCIP1, as well as in other TCIPs that were more potent in cells (for example see JWZ-7-6 or XFL-03-017, Ext. Data Fig. 2b,c). Indeed, relative potency in cells among the different TCIPs correlated with the total area under the TR-FRET curve, with TCIP1 giving the most robust killing among those tested (Ext. Data Fig. 2b).

Hypothesizing that TCIP1 may induce a molecular glue interaction important for potency, we carried out isothermal calorimetry (ITC) binary and ternary titrations of TCIP1, BRD4^BD1^, and BCL6^BTB^. Strikingly, both binary interactions of BCL6:TCIP1 BRD4:TCIP1 were weak (K_D_^BRD4:TCIP1^ = 5.08μM; K_D_^BCL6:TCIP1^> 1mM, Ext. Data Fig. 2d), but the ternary complex affinity was K_D_^BRD4:TCIP1:BCL6^ = 340 ±108 nM (mean ± s.d. of 3 replicates) (Fig. 2e, left). Using an orthogonal method, biolayer interferometry (BLI), we obtained a similar affinity of K_D_^BRD4:TCIP1:BCL6^ = 316 ±182 nM (Fig. 2e, right) and confirmed that there was little interaction between TCIP1 and BCL6^BTB^. We verified the published affinities of JQ1 to BRD4 and BI3812 to BCL6 using ITC and that the protein domains do not interact on their own (Ext. Data Fig. 2d). BLI measurements also revealed that the ternary complex has a slow off-rate of 0.026 seconds with a half-life of 30 seconds (Ext. Data Fig. 2e). Taken together, the data suggests TCIP1 induces a stable, cooperative protein-protein interaction between the bromodomain of BRD4 and the BTB domain of BCL6, that could be documented in future structural studies^13,48,49^.

Finally, to test if the ternary complex formed by TCIP1 can activate gene expression using endogenous levels of BCL6 and BRD4, we designed a BCL6 reporter from known BCL6 binding sites at promoters of cell death genes such as *TP53* and *CASP8*, based on ChIP-seq data in DLBCL cells, including the flanking 10 bp to capture any endogenous transcription factors co-binding (Fig. 2f). Transduction of KARPAS422 cells with the reporter and subsequent addition of TCIP1 revealed a dose-dependent activation with an EC_50_ of 4 nM, surprisingly similar to the EC_50_ of cell viability in these same cells (Fig. 2f). Reporter activation also featured the characteristic “hook effect” of a bifunctional molecule, with maximum activation occurring at 100 nM. We conclude from these experiments that activation of BCL6-dependent genes requires a ternary complex of BRD4, TCIP1 and BCL6.

### TCIP1 activates apoptosis at each stage of the cell cycle

Cancers can evade cell killing by many chemotherapeutics that function only during a specific stage of the cell cycle^50^. To characterize the rapid cell death observed with TCIP1, we quantified cells that have externalized phosphatidylserine by staining with annexin V. We observed a dose-dependent increase in annexin-positive cells, at 10 nM TCIP1, at 24hrs (Fig. 3a).

**Figure 3.**
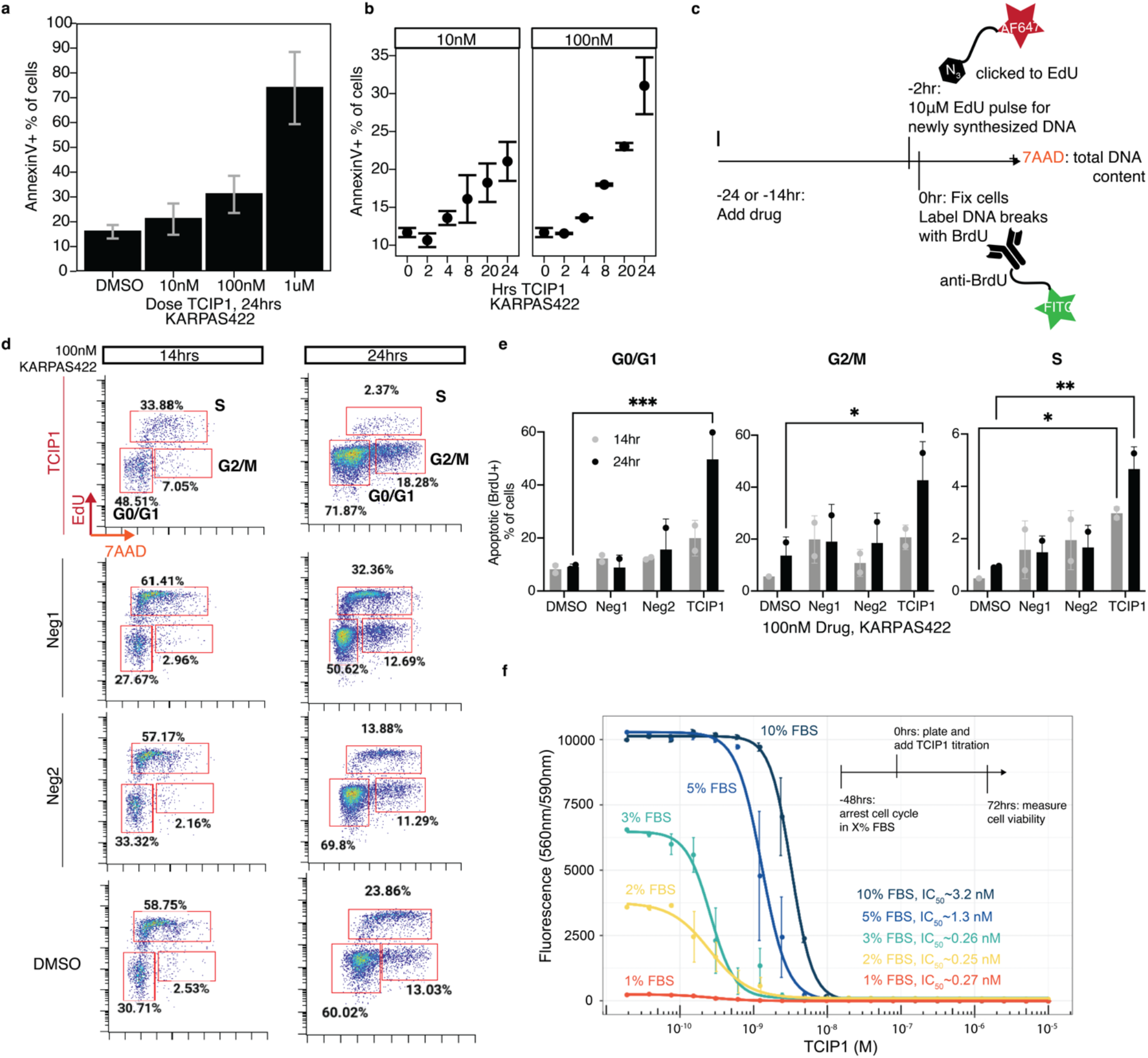
TCIP1 induces apoptosis at every stage of the cell cycle. **a.** Dose-dependent induction of apoptosis at 24hrs by TCIP1 as measured by AnnexinV-positive cells (n=4 biological repeats (6 for DMSO), mean±s.d.). **b.** Kinetics of TCIP1-induced apoptosis in KARPAS422 cells (n=3 biological repeats for 10 nM, 2 for 100 nM). **c.** Design of assay to measure cell cycle progression simultaneously with apoptosis using Terminal deoxynucleotidyl transferase dUTP nick end labeling (TUNEL) staining. **d.** TCIP1 induction of cell cycle arrest compared to controls. The cell cycle assay was repeated once. **e.** 100nM TCIP1 induction of apoptosis as measured by DNA fragmentation at each stage of the cell cycle (n=2 biological repeats, mean±s.d.). **f.** Measurement of cell viability after cell cycle arrest in G0/G1 by serum starvation in KARPAS422 cells and TCIP1 addition (n=3 biological repeats, mean±s.d.).

TCIP1induced detectable apoptosis by 4-8 hours (Fig. 3b), likely reflecting the fact that genes must be transcribed and translated for induction of cell death. While performing experiments, we observed that the doubling time of the KARPAS422 cell line is about 60-90 hours, yet killing was complete within 48 hours, suggesting that TCIP1-induced death is not cell cycle-dependent. To investigate this issue more carefully, we performed cell cycle analysis in concert with TUNEL staining (Fig. 3c) revealing that TCIP1 induced both a G1/S and G2/M block in the cell cycle (Fig. 3d). By examining DNA cleavage with the TUNEL assay, we found that cell death occurred during all phases of the cell cycle (Fig. 3e). The finding implies that unlike many conventional chemotherapeutics, BRD4-BCL6-TCIPs could kill cancer cells without periods of potential insensitivity. This observation is consistent with the total loss of cell viability produced by TCIP1 compared to JQ1 or BI3812 (Ext. Data Fig. 1a). To further examine the mechanism of cell death by TCIP1, we used serum starvation to arrest the cell cycle. Surprisingly, the cells became even more sensitive to TCIP1 after serum deprivation, exhibiting an EC_50_ of 250 pM compared to 1.1 nM without arrest (Fig. 3f). Taken together, these observations suggest that TCIP1 is an efficient inducer of apoptosis in lymphoma cells.

### TCIP1 reprograms gene expression

To define the genes involved in induction of apoptosis by TCIP1, we carried out RNA-seq studies 20 hours after adding drug at 10 or 100 nM, when apoptosis was just beginning and the critical genes were likely to be executing their function(s). Principal component analysis indicated that repeats were highly correlated and drug concentration determined most of the variation within the samples (Extended data Fig 4a). At 10 nM TCIP1, 1,674 genes were increased in expression, while 1,364 were reduced (Fig. 4a). Because a high percentage of the TCIP1-treated cells contain fragmented DNA by TUNEL assay at 24 hours (Fig 3e), many of the “reduced genes” could reflect DNA fragmentation as part of the apoptotic process. Among the group of genes whose expression was most reduced were MYC and its targets(Fig. 4b). We examined the top 100 most TCIP1-reduced genes using ChIP-seq data of human transcription factors in other cell lines^51^, and found the promoters of TCIP1-inhibited genes highly enriched for MYC binding in multiple datasets (Fig. 4c). This is likely significant since many DLBCLs are considered to be MYC-dependent^52,53^. Genes activated by TCIP1 were enriched for known BCL6 targets which are normally repressed by BCL6 (Fig. 4a and Ext. Data Fig. 4e)). For example, *CDKN1A* (p21) was activated by about 64-fold, while *FOXO3* and *PMAIP1/NOXA*, pro-apoptotic genes in lymphocytes, were also induced. Gene expression changes at 100 nM TCIP1 echoed those at 10 nM, albeit increased in magnitude (Ext. Data Figs. 4b,c). The changes in mRNA levels were paralleled by dose- and time-dependent changes in protein expression in two different DLBCL cell lines, SUDHL5 and KARPAS422 (Figs. 4d, e). JQ1 is known to reduce expression of MYC^38^, but at much higher levels of drug (500 nM) than TCIP1 (starting at less than 1 nM, within 2hrs at 10nM, see Figs. 4d,e), implying that the repression we see could not be due to JQ1 acting alone or synergistically with BI3812. In addition, p21, which is normally negatively regulated by BCL6, is induced by low nanomolar concentrations of TCIP1 (Fig. 4d). Of interest is the observation that FOXO3 is activated by 500 pM TCIP1 within 20 hours (Fig. 4d) and within two hours by 10 nM TCIP1 (Fig. 4e). FOXO3 is a master pro-apoptotic gene^54^ with a BCL6 binding site at its promoter^55^; it is normally activated downstream of TP53^56^. However, we note that TP53, a known BCL6 target often upstream of p21 and other pro-apoptotic genes, is biallelically inactivated in KARPAS422 and several of the other CHOP-resistant DLBCL that are robustly killed by TCIP1 (Ext. Data Fig. 1b). Additional experiments indicated that the chemical controls Neg1 and Neg2 do not affect TCIP1 targets (Ext. Data Fig. 5a). These studies indicate that TCIP1 effectively reprograms the transcriptional circuitry of DLBCLs such that cell killing is produced by a pathway that normally represses apoptosis.

**Figure 4.**
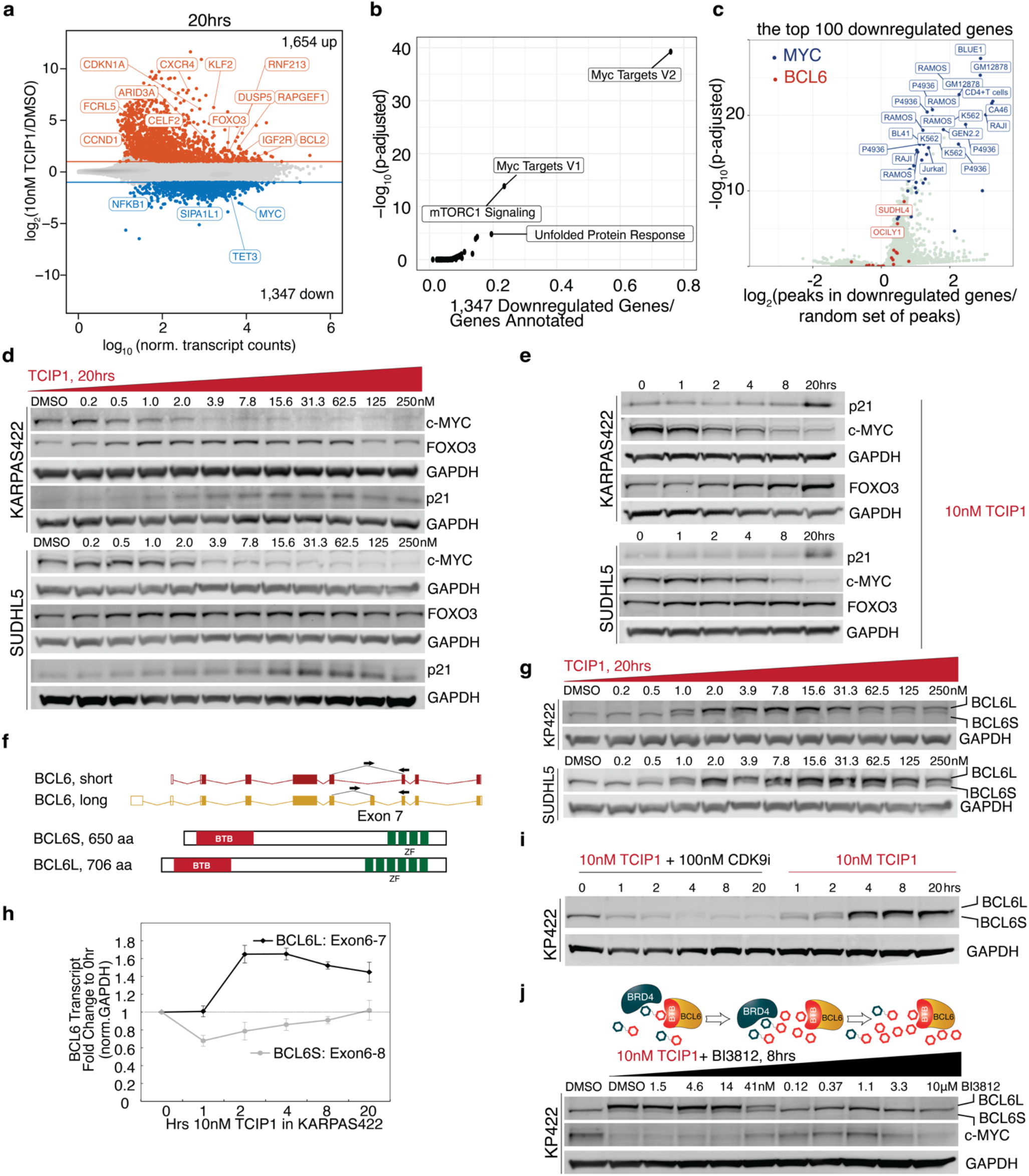
TCIP1 represses MYC target genes and rewires BCL6 auto-inhibition to a positive feedback pathway. **a.** Gene activation (median change: 4-fold up) and repression after 10 nM TCIP1 addition in KARPAS422 cells for 20hrs, with well-known BCL6 targets labeled. Significance cutoffs were p_adj_≤0.05 and |log_2_(Drug/DMSO)|≥1), n=2 biological repeats. **b**. Downregulated genes are significantly enriched for MYC targets (MSigDB Hallmark Pathways). **c**. Analysis of transcription factor binding at the top downregulated genes in over 3,000 public transcription factor ChIP-seq datasets from blood-lineage cells. **d**. Dose-dependent changes in protein levels of target genes selected from the RNA-seq results in two separate DLBCL cell lines, KARPAS422 and SUDHL5. **e**. Kinetics of protein changes in MYC, CDKN1A/p21, and FOXO3 in DLBCL cell lines after 10 nM TCIP1 treatment. **f.** BCL6 isoform structure. **g.** Induction of long isoform of BCL6 protein after treatment with as little as 0.5 picomolar TCIP1. **h**. mRNA levels measured by RT-qPCR of the long isoform and of the short isoform of BCL6, with primers specific to exon-exon junctions unique to each isoform (shown as arrows in **g**) n=3 biological repeats, mean±s.d. **i.** Simultaneous treatment of 10 nM TCIP1 and 100 nM CDK9 inhibitor, NVP2^60^, to inhibit transcriptional elongation and measurement of protein levels of BCL6 isoforms. **j.** Rescue of BCL6 upregulation and MYC downregulation by competitive titration of the BTB binder BI3812 against constant 10nM TCIP1 for 8hrs. All Western blots except **i** and **j** were repeated at least twice on independent passages of cell lines by 2 independent investigators.

### TCIP1 rewires the BCL6 negative feedback pathway to a positive feedback loop

BCL6 is overexpressed in several cancers either through mutation or other means^57^, apparently to protect the cancer cell from programmed cell death induced by TP53 and other pathways inhibited by BCL6. BCL6 expression is subject to negative autoregulation that originates from BCL6 binding sites in the 1^st^ intron of the BCL6 gene, which are often deleted or mutated in DLBCL^58,59^, providing protection from cell death. The TCIPs we have designed and synthesized should convert this negative feedback pathway to a positive feedback pathway by replacing epigenetic repression through HDACs^33^ and PRC^34^ with transcriptional activation by BRD4. To determine if this prediction is correct, we examined BCL6 levels after treatment with BRD4-BCL6 TCIP1 and found that BCL6 was significantly increased in a dose-dependent manner (Fig. 4g). This increase occurs within 1 hour (Fig. 4i) and was due to the long isoform of BCL6 (Fig. 4f), resulting from transcription originating from the 1^st^ exon induced by treatment with 10 nM TCIP1 (Fig. 4h). Upon simultaneous addition of TCIP1 and a nanomolar CDK9 inhibitor, NVP2^60^, to block elongation of transcription, BCL6 protein levels failed to be induced (Fig. 4i). Kinetics of BCL6 induction were similar in a second, TCIP1-sensitive DLBCL cell line, SUDHL5; in addition, BRD4 protein levels did not change substantially (Ext. Data Fig. 6b,c). The chemical controls Neg1 and Neg2 did not affect BCL6 (Ext. Data Fig. 6a).Because the primary transcript of BCL6 is 24 kilobases and RNA polymerase II moves at 2-3 kilobases per minute^61^, it could be transcribed, spliced and translated in 50 to 74 minutes. This indicates that transcription of the autoregulated long form of BCL6 starts almost immediately, less than 10 minutes after addition of TCIP1, and hence it is likely a direct target. Thus, the TCIP strategy turns a repressive negative feedback pathway into a positive feedback pathway, amplifying the potency of the molecule in killing cancer cells (model in Ext. Data Fig. 6d).

Finally, to clarify the role of ternary complex formation for the gene expression changes observed, we carried out a chemical competition experiment experiment by titrating the BCL6^BTB^ binder BI3812 against a constant 10 nM TCIP1 (Fig. 4j). Titration of BI3812 rescued both BCL6 upregulation and MYC downregulation after 8 hrs of drug addition (Fig. 4j). The results indicate that both activation and repression are linked to BRD4-TCIP1-BCL6 ternary complex formation.

### Identification of direct targets of TCIP1 by short timepoint RNA- and ChIP-seq

To tentatively identify direct targets of TCIP1 we carried out a short time-course RNA-seq with 10 nM TCIP1 in KARPAS422 cells for 1hr, 2hrs, and 4hrs along with negative controls (Fig. 5a,b). A selective set of only 140 genes, including well-characterized BCL6-repressed targets such as *BCL2L11/BIM^62^, FOXO3*, and *BCL6*, were activated at 2 hours and likely represent direct transcriptional targets of TCIP1 (Fig. 5a). Almost all differential genes were increasingly activated at 4 hours compared to negligible effects of the control molecules Neg1 and Neg2 (Fig. 5b). Consistent with protein kinetic data (Fig. 4e), *MYC* and its targets began to be repressed at 4 hours (Ext. Data Fig. 7a, c). Although a number of genes showed reduced expression in TCIP1 treated cells at these early times, we felt that they might be difficult to interpret because of the onset of DNA fragmentation. Analysis of BCL6 binding data using published BCL6 ChIP-seq in the DLBCL line OCILY1 and GC B Cells^55^ at the promoters of genes upregulated at 1, 2, and 4hrs showed that 53%, 57%, and 55% respectively of them had BCL6 enriched within 1 kilobase of their transcription start site (TSS) (Methods). Further analysis using ChIP-seq data of human transcription factors in other cell lines^51^, revealed that the promoters of these TCIP1-activated genes were statistically significantly enriched for BCL6 binding in multiple datasets (Fig. 5c). These studies indicate that TCIP1 is highly specific to activation of BCL6-target genes.

**Figure 5.**
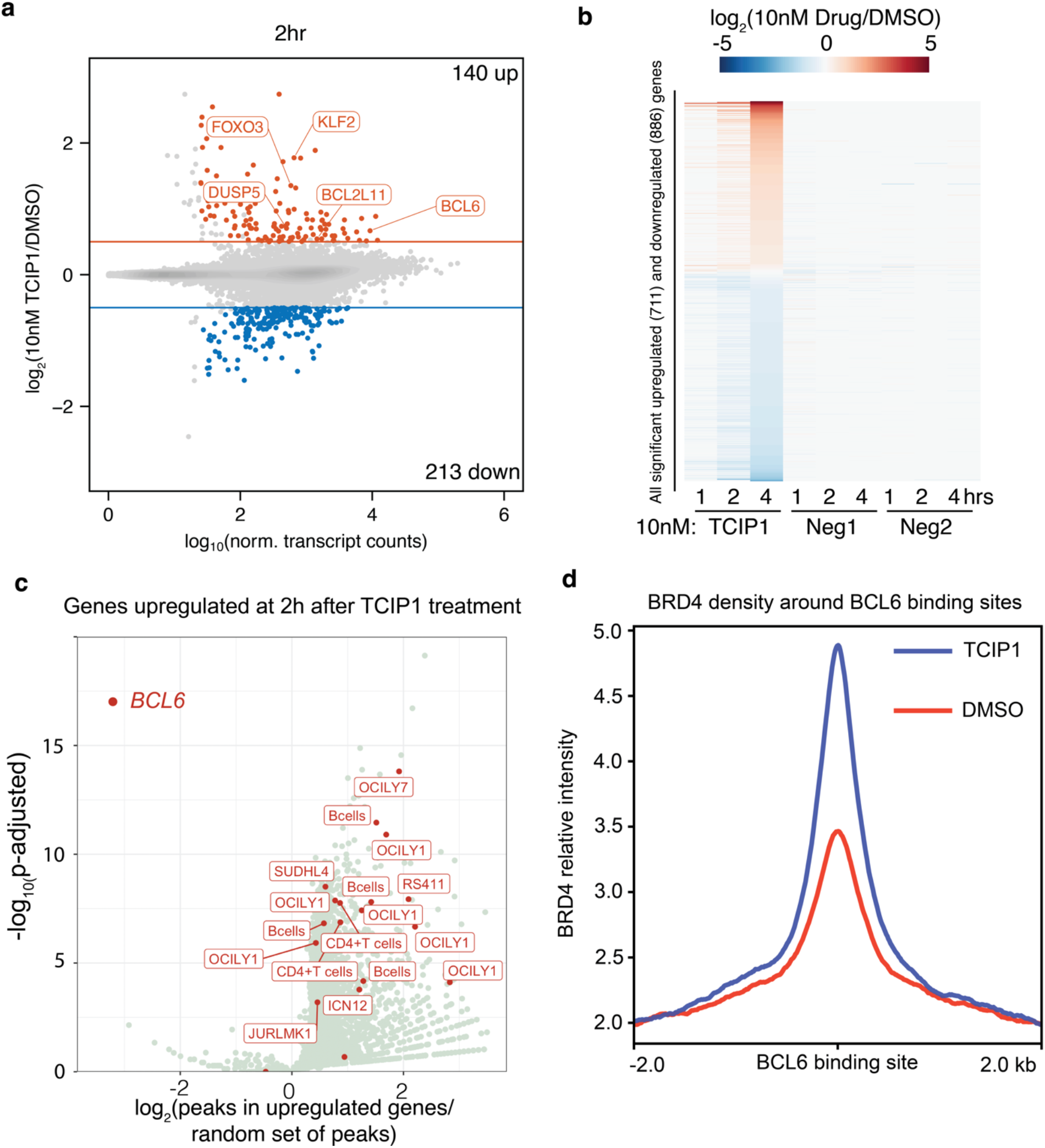
Rapid activation of BCL6 target genes by recruitment of BRD4. **a.** Gene expression changes by RNA-seq after 10 nM TCIP1 addition for 2hrs in KARPAS422 cells, with well-known BCL6 targets labeled. Significance cutoffs were p_adj_≤0.05 and |log_2_(Drug/DMSO)|≥0.5, n=3 biological repeats. **b.** Time-dependent changes in gene expression after 1,2, and 4hrs 10 nM TCIP1 compared to negligible effects of 10 nM Neg1 and Neg2 controls. **c.** Enrichment analysis of promoters of genes at 2hrs for transcription factor peaks in over 3,000 public transcription factor ChIP-seq datasets in blood-lineage cells. **d.** BRD4 ChIP-seq pekas in KARPAS422 cells at BCL6 summits after 1hr 100 nM TCIP1 addition.

ChIP-seq studies of BRD4 revealed that TCIP1 produced consistent, but modest (~1.5-fold) increase in BRD4 recruitment to BCL6 sites over the genome (Fig 5d). This was surprising because TCIP1 rapidly and robustly activated many genes normally repressed by BCL6. This observation could indicate that TCIP1 needs to recruit only small amounts of BRD4 to produce activation of BCL6 targets and/or that the antigenic determinant is disrupted by the extensive protein-protein interactions demonstrated by isothermal calorimetry within the ternary complex (Fig 1e).

### Generality of the TCIP strategy

We explored the generality and predictability of the TCIP approach by designing and synthesizing a series of molecules predicted to borrow the transcriptional activity of estrogen hormone receptor protein to activate derepressed BCL6 target genes and produce cell death (Fig. 6). We used the synthetic estrogen, estrone for these studies, which has somewhat lower affinity for the estrogen receptor (ER) than natural estradiol^63^ (Fig. 6a). Several of the ER-BCL6 TCIPs, such as TCIP2 (Fig. 6b), showed strong anti-proliferative activity with an EC_50_ of 355 nM (Fig. 6c). As predicted, killing was most robust in DLBCL lines such as KARPAS422 having higher expression of both ER and BCL6 (Fig. 6d). Interestingly, several ER-positive human breast cancer cells with low levels of BCL6 showed enhanced proliferation and survival, indicating that the estrone was active and that TCIPs are not intrinsically toxic (Fig 6d). In contrast, triple negative breast cancer cell lines with neither detectable BCL6 nor ER were not killed by the ER-BCL6 TCIP2 (Fig. 6d). These studies suggest that other transcriptional activators can be predictably hijacked or rewired to facilitate transcription of proapoptotic genes in DLBCL cells with high ER levels.

**Figure 6.**
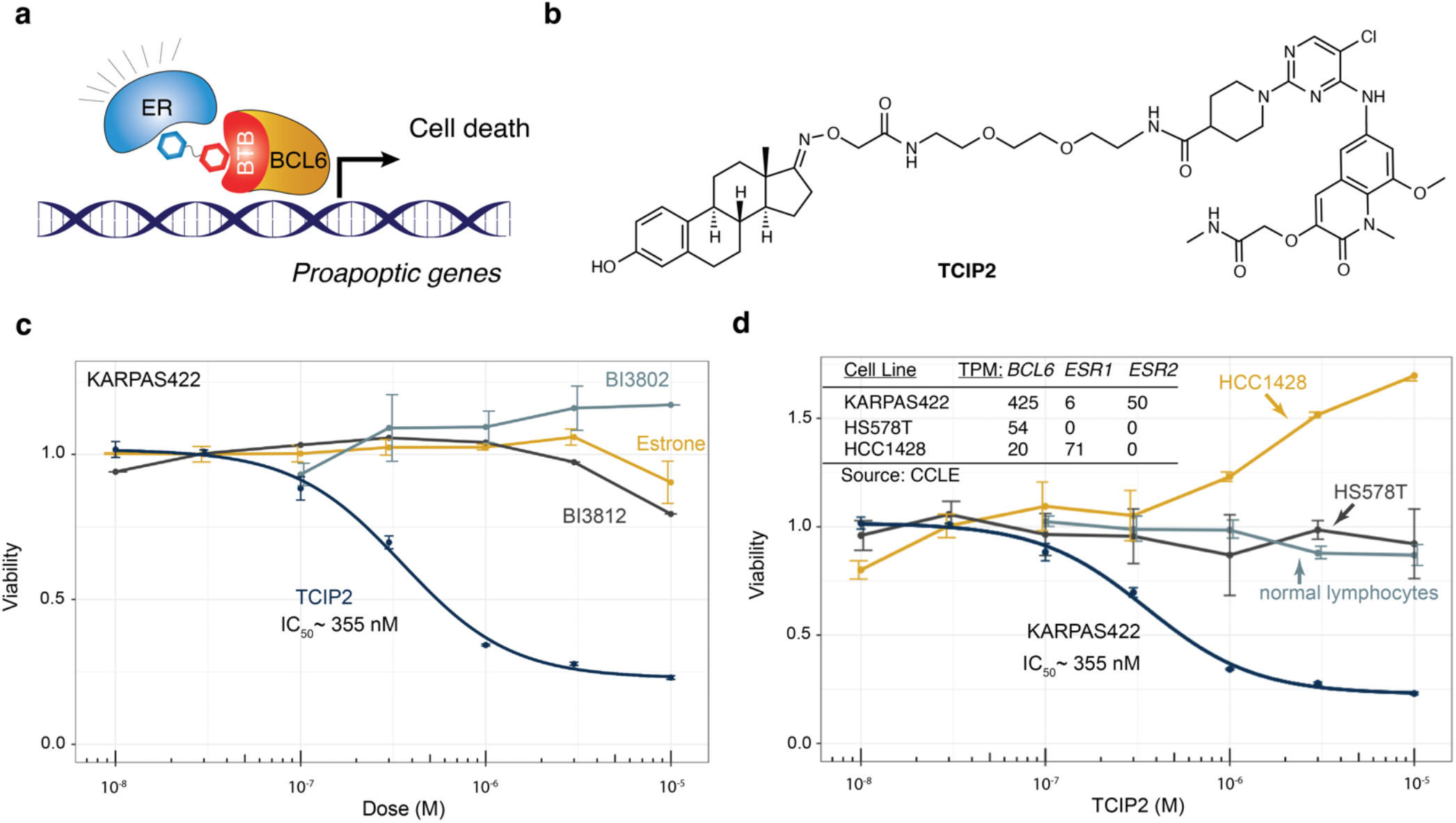
An estrone-BCL6 TCIP acts selectively in ER-positive cells. **a.** ER-BCL6 TCIP2 designed to induce cell death in estrogen-positive, BCL6-over-expressing DLBCLs. **b.** Chemical structure of TCIP2. **c.** Effect on cell viability of TCIP2 compared controls: estrone, BI3812, and BI3802 (BCL6 degrader) in KARPAS422 cells with high ER2626β *(ESR2)* levels, n=3 biological repeats, mean±s.d. **d.** Measurement of selective effect on cell viability by TCIP2 in DLBCL cells with coincident overexpression of ER and BCL6 (KARPAS422) compared to primary human lymphocytes, a triple-negative breast cancer cell line (HS578T), and ER-driven but BCL6-low breast cancer cells (HCC1428). n=3 biological repeats, mean±s.d.

### TCIP1 appears to be relatively non-toxic in primary human cells and mice

Cell death by TCIPs requires the coincident expression of both targets and hence, therapeutic effects might be more precisely targeted to the cancer cell rather than normal cells. To test this prediction, we examined primary human cells including fibroblasts and lymphocytes (Ext Data Fig. 8a,b, Fig 6d). T and B Lymphocytes are particularly germane because they have the highest level of BCL6^64^. While there is killing of both fibroblasts (EC_50_ ~ 470 nM) and lymphocytes (EC_50_ ~ 210 nM) by TCIP1 (Ext. Data Fig. 8), the levels required were at least 100-fold higher than that required for killing of DLBCL overexpressing BCL6, indicating the existence of a substantial therapeutic window. TCIP2, a less potent compound in DLBCL cell lines, had no effect in lymphocytes below 10 μM (Fig. 6d).

To set up future studies in tumor models, we evaluated the tolerability, pharmacokinetic properties, and target engagement of TCIP1 in wild-type mice, treating for 5 days with 10 mg/kg TCIP1 once daily by intraperitoneal injection. After treatment, mouse tissue was harvested for RNA-seq and histology. TCIP1 induced dramatic transcriptomic changes in the spleen (2,785 genes upregulated, 2,471 genes downregulated) compared to the liver and lung, despite comparable tissue concentrations of drug (Ext. Data Fig. 9a,b). Notable genes upregulated in DLBCL cells such as *FOXO3, CDKN1A*, or *HEXIM1* were also upregulated in the spleen as well as other known BCL6 targets in lymphocytes (Ext. Data Fig. 9c), suggesting engagement of BCL6 target genes. Despite the large transcriptomic changes in the spleen, TCIP1 was well-tolerated with no adverse effects noticed and no significant changes in mouse body weight (Ext. Data Fig. 9f). H&E staining also did not reveal noticeable abnormalities such as inflammatory infiltrates or apoptotic cells (Ext. Data Fig. 9g). The data supports the cellular evidence that TCIP1 acts in a context, tissue-specific manner dependent on coincident expression of BRD4 and BCL6. However, much additional toxicity testing in cell culture and in animal models must be performed before a conclusion can be reached as to their toxicity when used therapeutically.

## Discussion

While transcription factors are often not exquisitely specific in their pattern of expression, the outcome of transcriptional regulation often limits a protein to a specific cell type or biologic response. This specificity is thought to arise from the use of combinations of transcription factors that work together in enhancersomes to function as ANDGATES for the activation of different biologic processes^65,66^. One theoretical advantage of TCIPs is that they can exploit the fundamental mechanism used by transcription to produce specificity. Because TCIP activation of a specific gene requires coincident expression of two or more transcription factors, the approach might have greater specificity and less toxicity than simply inhibiting, degrading, or blocking production of a protein.

Existing approaches to cancer chemotherapy rely upon inhibiting or degrading a protein or preventing its synthesis by RNAi or CRISPR(i). These approaches require complete or near complete removal of the driver function. However, TCIPs rewire these drivers to cell death pathways, and need only to use a fraction of the total driver molecules per cell. This assertion is suggested by the fact that 10 nM TCIP1 produces only a modest, ~1.5-fold increase in BRD4 at BCL6 sites over the genome (Fig 5), despite robust gene activation and cell killing. This observation could also explain the far more robust cell killing seen with substantially lower concentrations of TCIP1, compared to the weaker anti-proliferative effects of conventional small molecule inhibitors or degraders of BCL6^35,67–71^ or BRD4^37^, which act by “occupancy-driven” pharmacology requiring binding to almost all copies of their cognate protein. As a consequence, TCIPs may avoid mechanism-based toxicity that almost inevitably occurs when one reduces the expression or activity of an essential protein, like BRD4.

Genomic studies demonstrate that cancers often have more than a single driver that may independently contribute to proliferation, survival, invasion, and metastasis and thereby complicate therapeutic approaches.^5,72^. In future versions, TCIPs can be designed to take advantage of any of these diverse biologic mechanisms that are required for the complex cancerous phenotype, requiring only chemical binders to the drivers of interest. By making use of the cancer cell’s own intrinsic driving pathways and rewiring them to activate pathways of cell death, we have introduced a new concept in cancer chemotherapy that is analogous to a dominant, gain-of-function in genetics. The TCIP strategy also engages genomic information specific to the tumor and its stage of development. In conventional chemotherapy multiple drivers are a distinct disadvantage and can produce resistance to single agents^5^. In contrast, the dominant gain-of-function produced by TCIPs could allow the killing of cancer cells with any number of independent drivers. Resistance to treatment often reflects the existence of multiple drivers or emerges when a second driving mechanism become active. Because of the potential versatility of the TCIP approach, an agile response to the emergence of resistance could be envisioned by design of additional TCIPs to the new drivers.

Past studies have used chemically induced proximity of genetically modified transcription factors or epigenetic regulators to activate or repress signal transduction or transcription of exogenous or endogenous genes^13,22,24^. While these studies were mechanistically informative and provided a catalogue of the biologic processes regulated by CIP^25^, they had little therapeutic potential because of the need to introduce genetically modified transcription factors or epigenetic regulators. Our experiments developing TCIPs to activate reporters and endogenous genes in unmodified cells suggest that many other chromatin modifiers and transcriptional regulators may be susceptible to chemical induced proximity-based control. The activation of endogenous genes by small molecule TCIPs might have application to many other areas of biology and medicine.

For example, TCIPs could be designed for use in activating death pathways in senescent cells, activating expression of therapeutic or haploinsufficent genes, activating the expression of neoantigens in human immunotherapy, or regulating gene expression in cells/organisms for synthetic biology applications. The fact that TCIPs take advantage of the mechanism of transcription specificity – combinatorics – might allow one to engineer exquisite specificity and hence low toxicity into these many uses.

## Author Contributions

G.R.C. conceived the TCIP strategy to regulate endogenous genes and of the rewiring cancer drivers to activate cell death pathways. S.G. contributed many inventive ideas and performed many of the studies shown in the figures was well as designed the first effective TCIP. A.K. contributed many inventive ideas to the underlying concept, defined the conditions for TCIP use and carried out the first successful TCIP experiments. C-Y.C. suggested the use of BCL6 as a means of producing cell death. S.H.K. screened the cell death genes to detect those that would directly kill cancer cells by over expression. N.S.G. and S.G. designed the first effective TCIPs. X.L. synthesized the first effective TCIP, and T.Z. contributed to TCIP design and optimization. W.J. synthesized TCIP1 (JWZ-7-7), the most potent TCIP to date. W.W. contributed innovative ideas to the computational analysis for the selection of TCIP components and helped fund the studies by writing grants with G.R.C. J.M.S. carried out experiments designed by G.R.C., S.G. and A.K. The manuscript was written by G.R.C, S.G., A.K., N.S.G. with input from all authors.

## Acknowledgements

The studies described in this manuscript were funded from a grant from HHMI to GRC, NIH grant CA 163915 and NIH grant 1 R01 MH126720-01 to GRC. Funding was also provided by a grant from the Mary Kay Foundation and a grant from the Stanford SPARK program. Funding was provided to SG from NIH grant 5F31HD103339-03. Funding was provided to NSG from departmental funds from Chemical and Systems Biology and the Stanford Cancer Institute and the Gray lab also receives or has received research funding from Novartis, Takeda, Astellas, Taiho, Jansen, Kinogen, Arbella, Deerfield, Springworks, Interline and Sanofi. Funding for PK studies was provided by NIH grant number 1 S10OD030332-01. S.G. would like to thank T. Reindl, E. Bruguera, and S. Hinshaw for helpful advice in biochemical studies. All authors would like to thank members of the Crabtree and Gray laboratories for feedback.

## Methods

### Cell Culture

Lymphoma cells were cultured in RPMI-1640 (ATCC 30-2001) + 10% FBS with antibiotics (100X PenStrep GIBCO #15140122). Primary human fibroblasts were obtained from ATCC (#CRL2522), cultured in DMEM+10% FBS with antibiotics and used at passages 3-5. Primary human tonsillar lymphocytes were a kind gift from M. M. Davis. Cells were routinely checked for mycoplasma and immediately checked upon suspicion. No cultures tested positive.

### Cell Viability Measurements

30,000 cells were plated in 100 μL media per well of a 96 well plate and treated with drug for indicated times and doses. A resazurin-based indicator of cell health (PrestoBlue, ThermoFisher #P50200) was added for 1.5hrs after which the fluorescence ratio at 560/590nm was recorded. The background fluorescence was subtracted and the signal was normalized to DMSO-treated cells. EC_50_ measurements on cell lines were done with 4 biological repeats by separate cell passages maintained by 3 independent investigators. Fit of dose-response curves to data and statistical analysis was performed using drc package.

### PRISM Cell Proliferation Assay

The PRISM cell proliferation assay was carried out as previously described^46^.

### Chemical Synthesis

Additional details are provided in the Supplementary Methods section.

### Protein Expression and Purification

The construct for 6xHis-TEV-BRD4_BD1 was described in Filippakopoulos, Qi et al^37^ and was a gift from Nicola Burgess-Brown (Addgene plasmid # 38943; http://n2t.net/addgene:38943; RRID:Addgene_38943). The construct for BCL6_BTB-AviTag was based off previously designed BCL6 constructs used for TR-FRET assays, as reported in multiple papers including^35,36^ and contains amnio acids 5-129 with three mutations, C8Q, C67R, C84N, that enhance stability but have no difference on backbone structure with the wild-type version^73^. A Trx-6xHis-HRV3C-BCL6_BTB construct without the AviTag was produced similarly for ITC studies where the tag was cleaved by addition of HRV3C. Additional details are provided in the Supplementary Methods section.

### TR-FRET

Each reaction contained 100 nM BRD4^BD1^, 100 nM BCL6^BTB^-Avi-Biot, 20nM Streptavidin-FITC (Thermo #SA1001), and 1:400 anti-6xHis terbium antibody (PerkinElmer #61HI2TLF) in 10μL of buffer containing 20mM HEPES, 150mM NaCl, 0.1% BSA, 0.1% NP-40, and 1mM TCEP in a 384well plate. Protein was incubated with drug digitally dispensed (Tecan D300e) for 1hr in the dark at room temperature before excitation at 337nm and measurement of emission at 520nm (FITC) and 490nm (terbium) with a PHERAstar FS plate reader (BMG Labtech). The ratio of signal at 520nm to 490nm was calculated and normalized to DMSO-treated conditions, and plotted.

### Isothermal Calorimetry (ITC)

Tag-cleaved version of BCL6^BTB^ and BRD4^BD1^ were used for experiments, in a VP-ITC machine. For binary assays with TCIP1, 400μM BCL6^BTB^ or BRD4^BD1^ were titrated from the syringe into a cell containing 40μM TCIP1. For the binary protein-protein ITC, 330μM BCL6^BTB^ was titrated into 68μM BRD4^BD1^. For binary assays with JQ1 or BI3812, 100μM BCL6^BTB^ or 350μM BRD4^BD1^ was titrated from the syringe into a cell containing 5μM BI3812 or 20μM JQ1. For the ternary complex assays, 200μM BRD4^BD1^ was incubated with 10μM TCIP1 in the cell (20-fold excess, to drive saturation of the binary complex), and 100μM BCL6^BTB^ was titrated from the syringe, at 310 rpm stirring at 25 ^O^C in a buffer containing 10mM HEPES pH 7.5, 200mM NaCl, 5% glycerol, 1mM TCEP, and matched DMSO % (never more than 0.4%) in the syringe and the cell. Data was fit to a one-site model using MicroCal LLC Origin software.

### Biolayer Interferometry (BLI)

The tag-cleaved version of BRD4^BD1^ and biotinylated BCL6^BTB^ were used for experiments, in a Gator Bio BLI machine. 50μM of BRD4^BD1^ was added to each well containing titrations of TCIP1 from 5.5nM to 12μM so that BRD4^BD1^ would be in excess and drive binary BRD4^BD1^:TCIP1 complex formation. 100nM BCL6^BTB^ was loaded on the tip. Experiments were carried out at 25 ^O^C. After loading, association was carried out for 300s, dissociation for 300s, and a baseline for 30s. A TCIP1-only control was carried out for each concentration confirming that there was no binding between BCL6^BTB^ and TCIP1 on its own. A BRD4^BD1^-only control was carried out similarly confirming that BCL6^BTB^ and BRD4^BD1^ do not interact on their own. Data was analyzed in GraphPad Prism with the association curves fit to the model “One-phase association” and the dissociation curves to the model “One-phase decay” to obtain kinetic parameters. The K_D_ was obtained by fitting a “One-site binding” curve to the Span of each association curve vs. concentration of drug.

### Flow Cytometry

For annexin V assays, 500,000 cells plated at 1M/mL treated with drug for indicated timepoints and doses were harvested on ice and washed twice in 1% BSA/PBS. 2.5 μL of 7-AAD and 2.5 μL of FITC-Annexin V (Biolegend #640922) were added. Cells were incubated for 15mins at RT, then immediately measured on a BD Accuri. Gates were drawn based off single-stain and no-stain controls.

For cell cycle and TUNEL analysis, cells plated at 1M/mL were treated with drug for indicated timepoints and doses and pulsed with 10μM ethynyl-EdU (Thermo #C10424) for 2hrs prior to harvesting on ice. 1M cells were counted and washed in 1% BSA/PBS. Cells were resuspended at 10M/mL and fixed in 4% PFA, washed, and permeabilized in 0.5% Triton-X100/PBS. Fixed and permeabilized cells were washed and labeled with BrdUTP using terminal deoxynucleotidyl transferase (TdT, BD #556405) for 60mins at 37 ^O^C, rinsed, and then labeled with AlexaFluor 647-azide (Thermo #C10424) for 30mins at RT in the dark. After washes, the sample was incubated with 2μL 7-AAD and 5μL RNAseA for 30mins at RT in the dark, washed, and measured on a BD Accuri. Gates were drawn based off single-stain and no-stain controls and kept constant across conditions.

### BCL6 Reporter Assay

KARPAS cells were lentivirally transduced construct containing the reporter. After selection, cells were plated and treated with indicated amount of TCIP1 for 8hrs. Cells were washed in 1% FBS/PBS, 2.5 μL of 7-AAD was added to distinguish live from dead cells, and harvested for flow cytometry on a BD Accuri. Gates were drawn off non-transduced cells and transduced cells treated with DMSO.

### RNA Extraction, qPCR, and Sequencing Library Preparation

Cells were plated at 1M/mL and harvested in TRIsure (Bioline #38033). RNA was extracted using Direct-zol RNA MicroPrep columns (Zymo #R2062) treated with DNAseI. cDNA was prepared for RT-qPCR using the SensiFAST cDNA preparation kit according to manufacturer instructions (Bioline #65054). 1μL of cDNA was used per RT-qPCR reaction prepared with SYBR Lo-ROX (Bioline #94020). For sequencing library preparation, polyA-containing transcripts were enriched for (NEB #E7490S) and prepared into paired-end libraries (NEB #E7760S). Libraries were sequenced on an Illumina NovaSeq (Novogene).

### Western blots

Cells were plated at 1M/mL and treated with drug at indicated timepoints and doses. 2M Cells were harvested on ice in RIPA buffer (50mM Tris-HCl pH 8, 150mM NaCl, 1% NP-40, 0.1% DOC, 1% SDS, protease inhibitor cocktail (homemade), 1mM DTT) and 1:200 benzonase (Sigma #E1014) was added and incubated for 20mins. After 10min centrifugation at 14,000g and 4 ^O^C, the supernatant was collected and protein concentration was measured by Bradford. Antibodies used for immunoblots are: BCL6 (Cell Signaling #D65C10), BRD4 (Abcam #ab243862), BCL2 (Cell Signaling #15071), TP53 (Santa Cruz DO-1), c-MYC (Cell Signaling D84C12), FOXO3 (Cell Signaling 75D8), p21 (Cell Signaling 12D1), GAPDH (Santa Cruz 6C5).

### RNA-seq Analysis

Raw reads were checked for quality using fastqc (https://www.bioinformatics.babraham.ac.uk/projects/fastqc/) and trimmed from adapters using cutadapt^74^ using parameters cutadapt-a AGATCGGAAGAGCACACGTCTGAACTCCAGTCA-b AGATCGGAAGAGCGTCGTGTAGGGAAAGAGTGT --nextseq-trim=20 --minimum-length 1. Transcripts were quantified using kallisto^75^ against the human Gencode v33 indexed transcriptome and annotations. Differential gene analysis was performed using DESeq2^76^ and pathway and enrichment analyses using Enrichr^77^ and ChIP-Atlas^51^. For analysis of BCL6 binding at at ±1kb from the TSS of differentially regulated genes, BCL6 peaks were reconstructed from OCILY1 DLBCL cells as deposited in^55^, using macs2^78^ callpeak with a stringent score cutoff >=100 and overlap was calculated.

### ChIP-seq Experiment and Library Preparation

30 million cells were treated with TCIP1 or DMSO for 1h. Cells were washed in PBS and crosslinked for 12 min in CiA FiX Buffer (50mM HEPES pH 8.0, 1mM EDTA, 0.5mM EGTA, 100 mM NaCl) with addition of formaldehyde to a final concentration of 1%. The crosslinking reaction was quenched by glycine added at 0.125M final concentration. Crosslinked cells were centrifuged at 1,000 x g for 5min. Nuclei were prepared by 10 min incubation of resuspended pellet in CiA NP-Rinse 1 buffer (50mM HEPES pH 8.0, 140mM NaCl, 1mM EDTA, 10% glycerol, 0.5% IGEPAL CA-630, 0.25% Triton X100) followed by wash in CiA NP-Rinse 2 buffer (10mM Tris pH 8.0, 1mM EDTA, 0.5 mM EGTA, 200mM NaCl). The pellet was resuspended in CiA Covaris Shearing Buffer (0.1% SDS.1mM EDTA pH 8.0, 10 mM Tris HCl pH 8.0) with 1000x protease inhibitors (Roche) and sonicated for 20 min with Covaris E220 sonicator (Peak Power 140, Duty Factor 5.0, Cycles/Burst 200). The distribution of fragments was confirmed with agarose gel. 300 μL of chromatin per ChIP was used with anti-BRD4 antibodies (BRD4 (E2A7X) Rabbit mAb #13440). The sequencing library preparation was performed using NEBNext Ultra II DNA kit (#E7645S). Libraries were sequenced with Illumina NovaSeq at Novogene.

### ChIP-seq Analysis

The data quality was checked using fastqc. The raw reads were trimmed from adapters with trim_galore (parameters: --paired–illumina) and raw reads were aligned to hg38 human genome assembly using bowtie2 (parameters: --local). Low quality reads, duplicated reads and reads with multiple alignments were removed using samtools^79^ and Picard (https://broadinstitute.github.io/picard/). Macs2^78^ was used to map position of BRD4 peaks. Bedtools^80^ was used to find consensus set of peaks by merging peaks across multiple conditions (bedtools merge), count number of reads in peaks (bedtools intersect-c) and generate genome coverage (bedtools genomecov-bga). deepTools^81^ was used to generate BRD4 coverage densities across multiple experimental conditions. The peak differential analysis was performed using DESeq2^76^. The SRX4609168 public dataset was used to extract positions of BCL6 peaks.

### Data Reporting

No statistical methods were used to predetermine sample size. Experiments were not randomized and investigators were not blinded.

### Data Availability

Uncropped blots are available in Supplementary Figure 1. Coomassie gels of recombinant proteins are available in Supplementary Figure 2. Flow gating strategy is available in Supplementary Figure 3. Sequencing data has been deposited to GSE211282. Reagents generated in this study will be made available upon request with a completed Materials Transfer Agreement. Other data and materials available from authors on request.

---

**Extended Data Figure 1.**
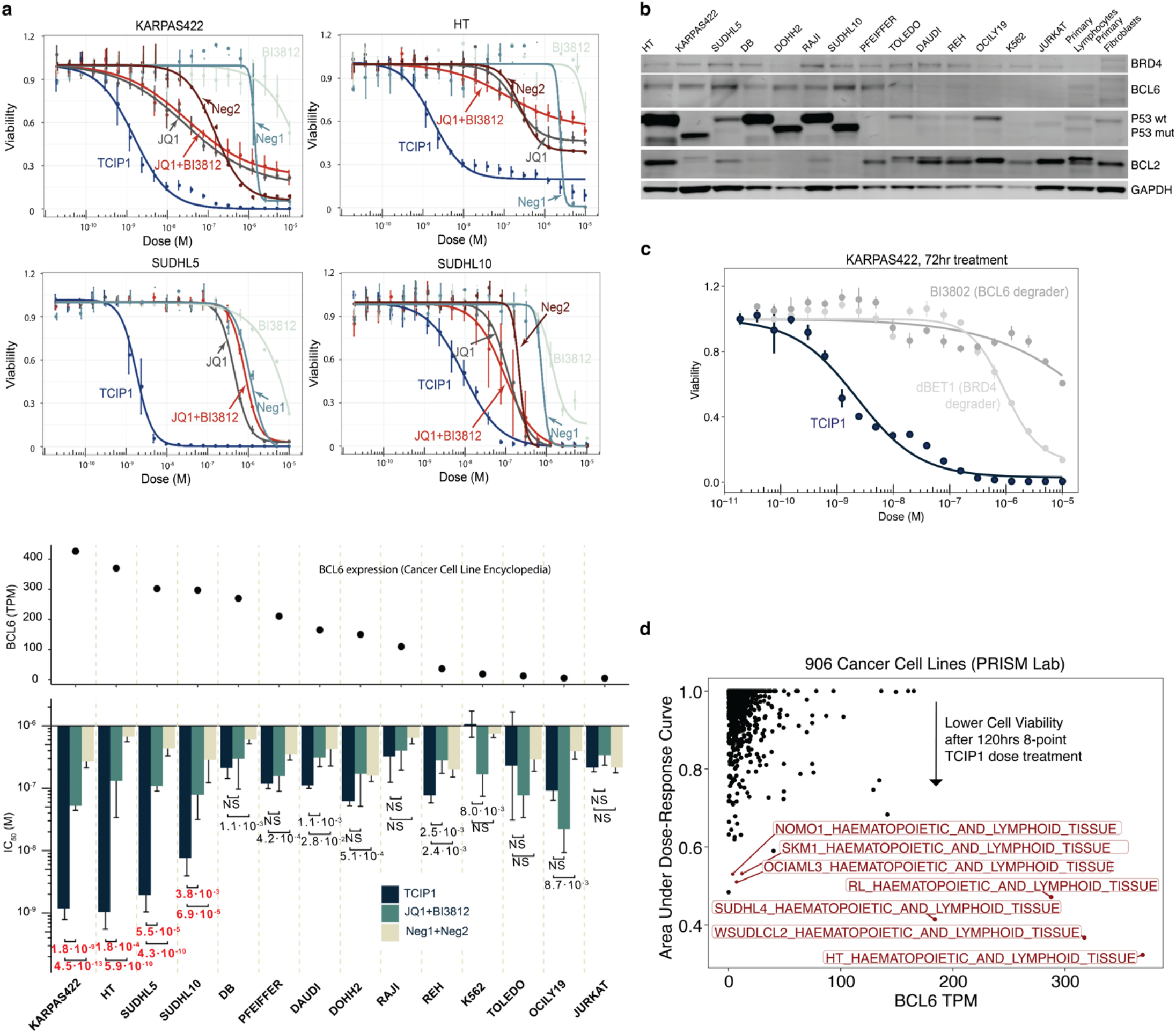
TCIP1 is selective to BCL6-expressing cancer cell lines compared to controls. **a.** Comparison of TCIP1 effect on cell viability to effect of negative controls Neg1 and Neg2, or single-sided molecules JQ1 and BI3812, or the additive effect of JQ1+BI3812. TCIP1 EC_50_ of cell viability is anti-correlated with BCL6 content across 14 different cancer cell lines, n=3 biological repeats, mean±s.d. **b.** Measurement of P53 or BCL2 status of DLBCL cell lines ranked from left to right from high to low-BCL6 protein content. **c.** Comparison of TCIP1 effect on cell viability to effect of BRD4 or BCL6 degraders (n=3 technical repeats). **d.** Unbiased screen of the effect of TCIP1 on the viability of 906 barcoded cancer cell lines (PRISM). Drug was dosed for 120hrs in triplicate (Methods).

**Extended Data Figure 2.**
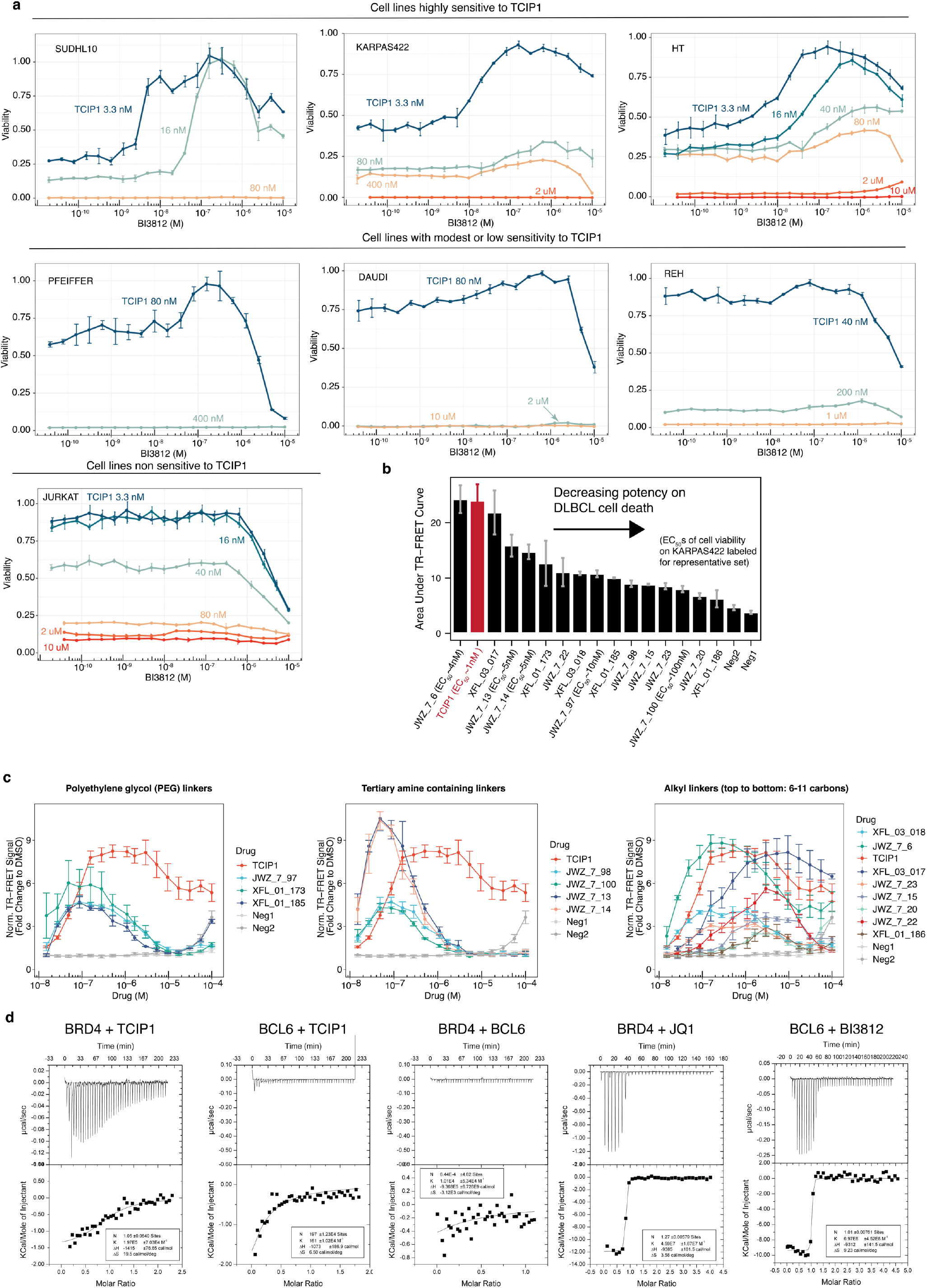
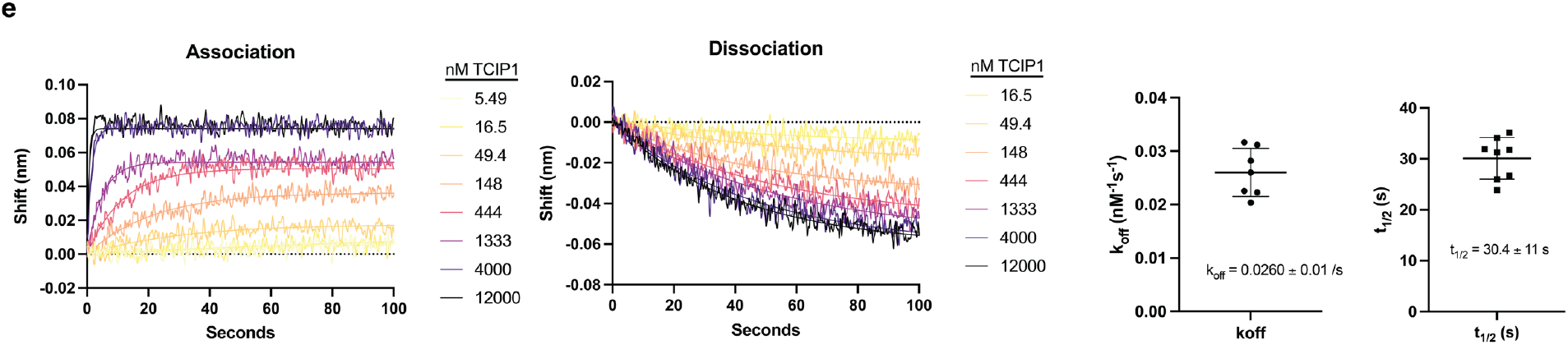
Ternary complex formation in cells and *in vitro*. **a.** Rescue of TCIP1-induced cell death across cancer cell lines that are highly sensitive to TCIP1, moderately sensitive, or not at all sensitive. n=3 biological repeats, mean±s.d.. **b.** Area under curve of TR-FRET correlates with potency of TCIP on cell death (KARPAS422 cells, viability at 72hrs). Representative cellular EC_50_s labeled. **c.** Ternary complex formation by TCIPs with related chemistries. TCIP1 and Neg1 and Neg2 controls are plotted on every graph. n=3 repeats, mean±s.e. **d.** Isothermal calorimetry experiments to measure binary affinities of TCIP1 to BRD4 (BD1), BCL6 (BTB), and associated controls. Representative data from n=2 shown. **e.** Representative biolayer interferometry measurements (BLI) of ternary complex kinetics from n=2 repeats shown with biotinylated BCL6 (BTB) on the tip and excess BRD4 in the well with titration of TCIP1.

**Extended Data Figure 3.**
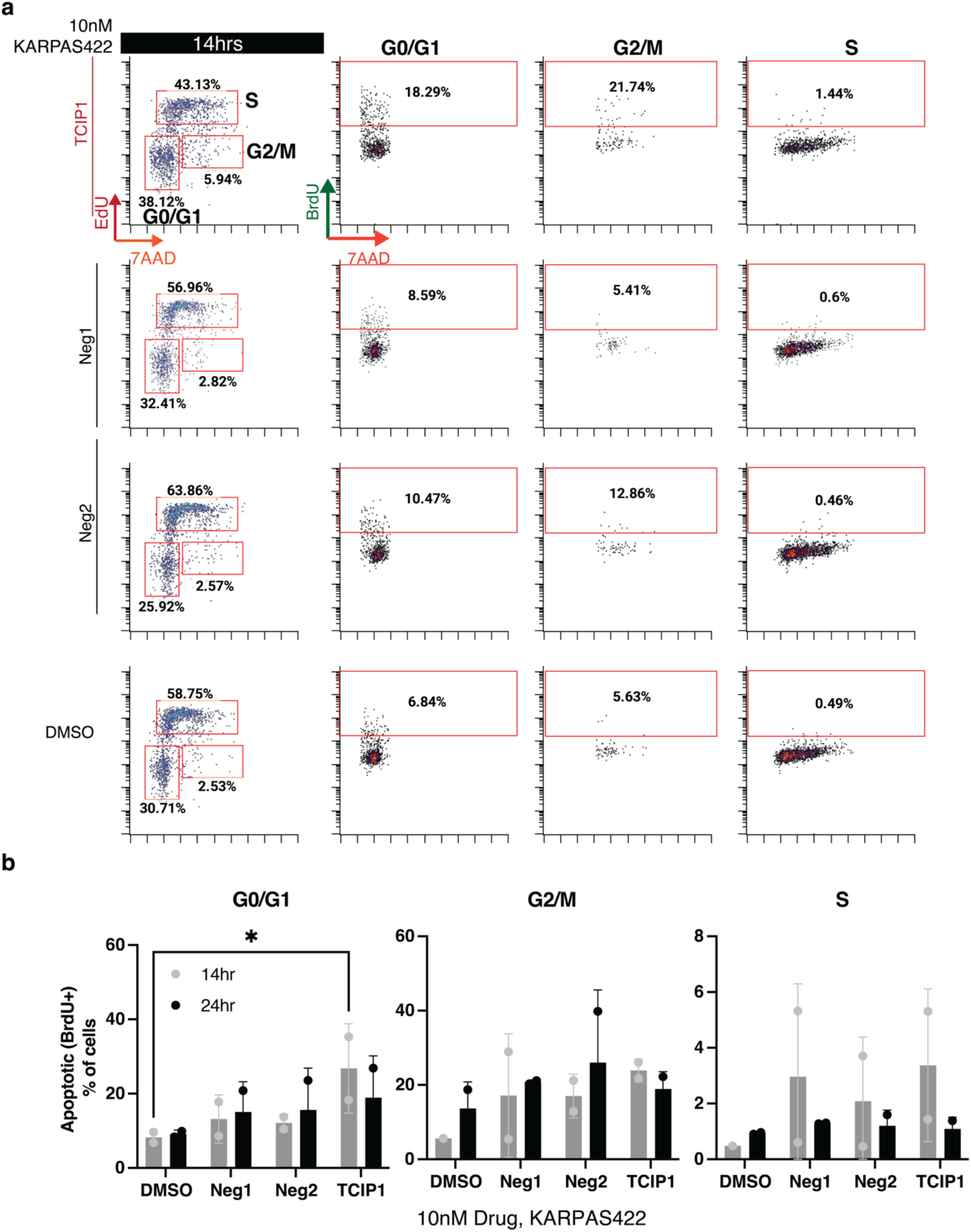
Early detection at 10 nM TCIP1 of cell cycle block and apoptosis. **a.** 10 nM TCIP1 addition at 14hrs and measurement of cell cycle block and apoptosis in KARPAS422 cells. **b.** 10nM TCIP1 induction of apoptosis as measured by DNA fragmentation at each stage of the cell cycle (n=2 biological repeats, mean±s.d.). (n=2 biological repeats, mean±s.d.)

**Extended Data Figure 4.**
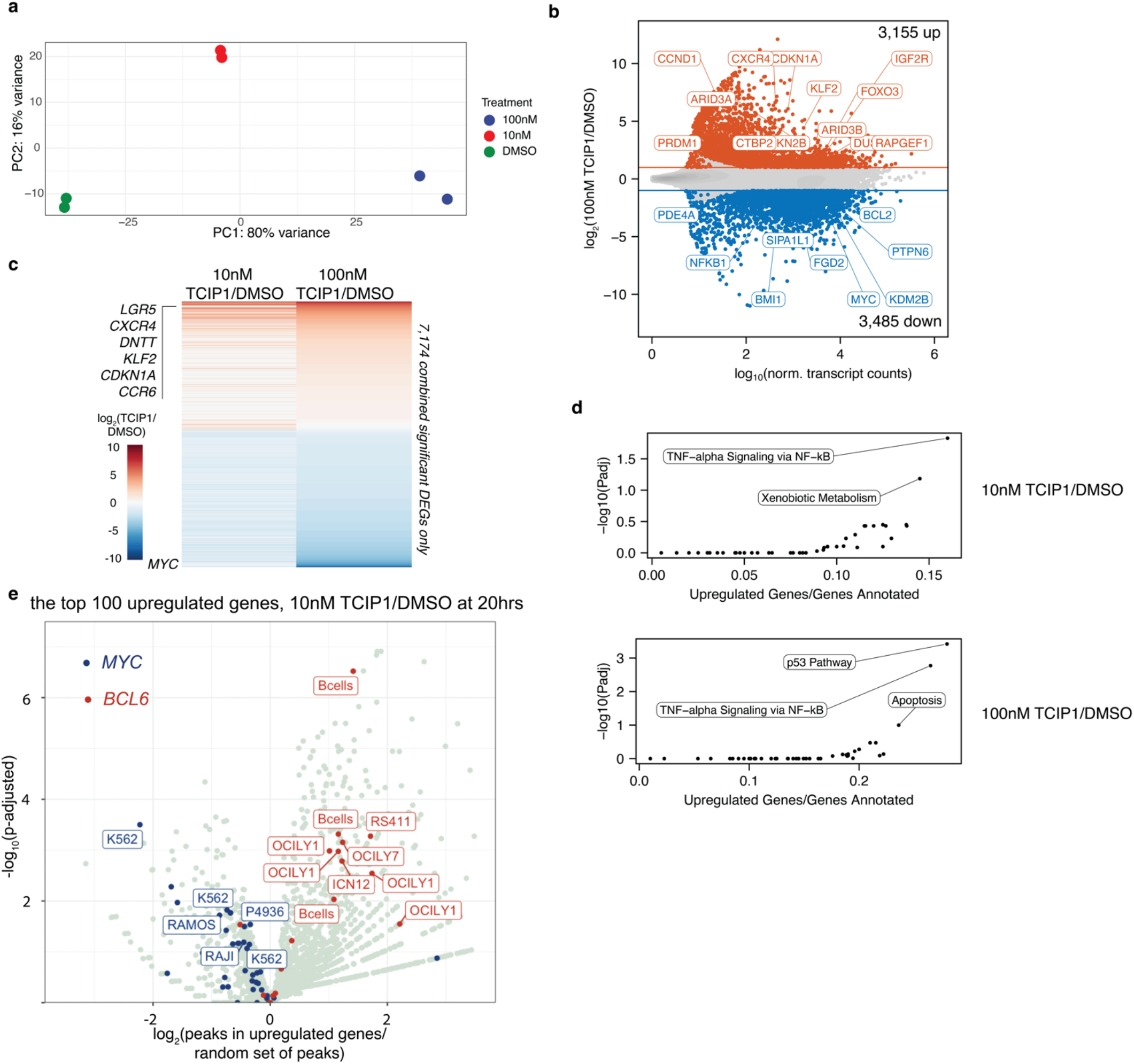
Robust and dose-dependent gene regulation by TCIP1. **a.** Principal component analysis of RNA-seq data after addition of TCIP1 for 20hrs in 2 biological replicates of KARPAS422 cells. **b.** Gene expression changes after addition of 100 nM TCIP1 for 20hrs in KARPAS422 cells. Significance cutoffs were p_adj_≤0.05 and |log_2_(Drug/DMSO)|≥1), n=2 biological repeats. **c.** Dose-dependent change in gene expression. **d.** Enrichment analysis of upregulated genes (MSigDB Hallmark Pathways). **e.** Analysis of TF binding at the top upregulated genes in over 3,000 public transcription factor ChIP-seq datasets from blood-lineage cells.

**Extended Data Figure 5.**
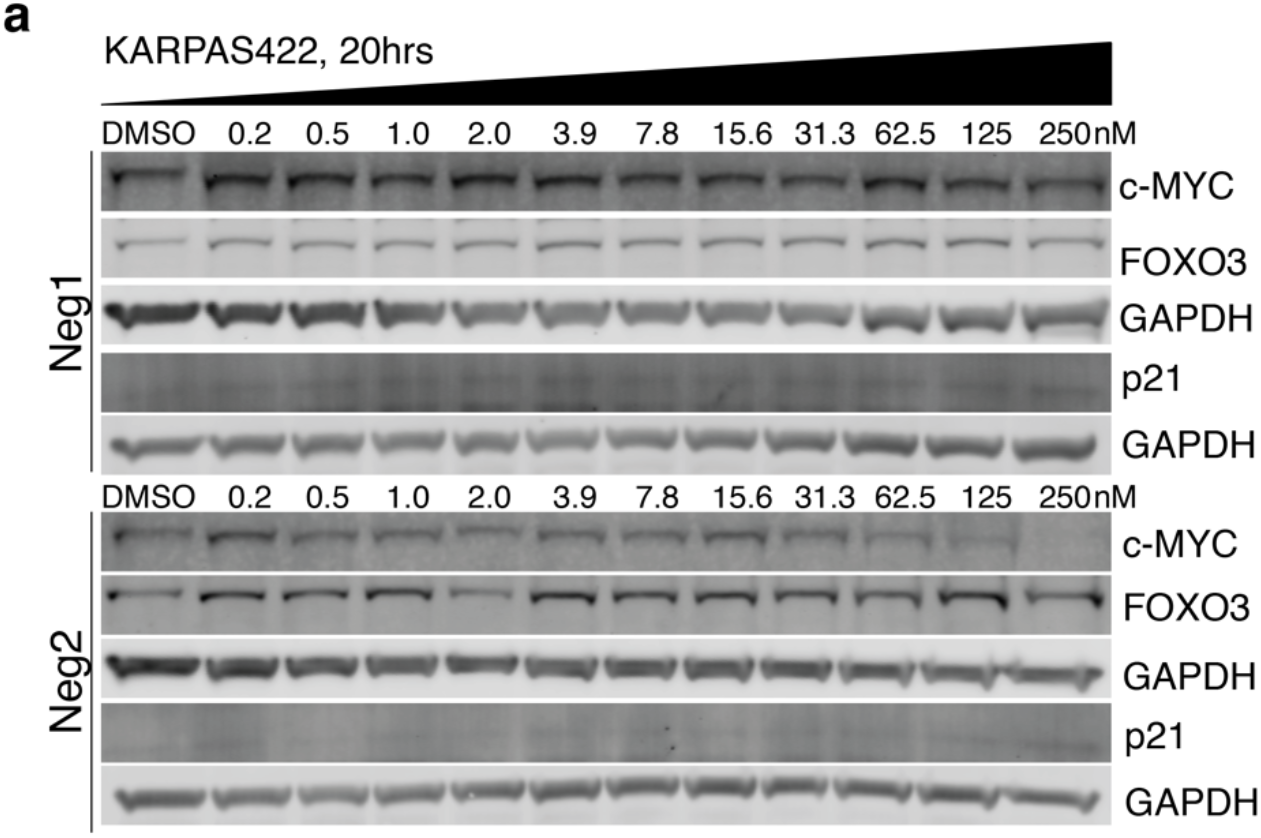
Negative controls do not affect TCIP1 targets. **a.** Dose-dependent changes in protein levels of target genes selected from the RNA-seq results in KARPAS422 at 20hrs after treatment with Neg1 or Neg2 (representative of n=2).

**Extended Data Figure 6.**
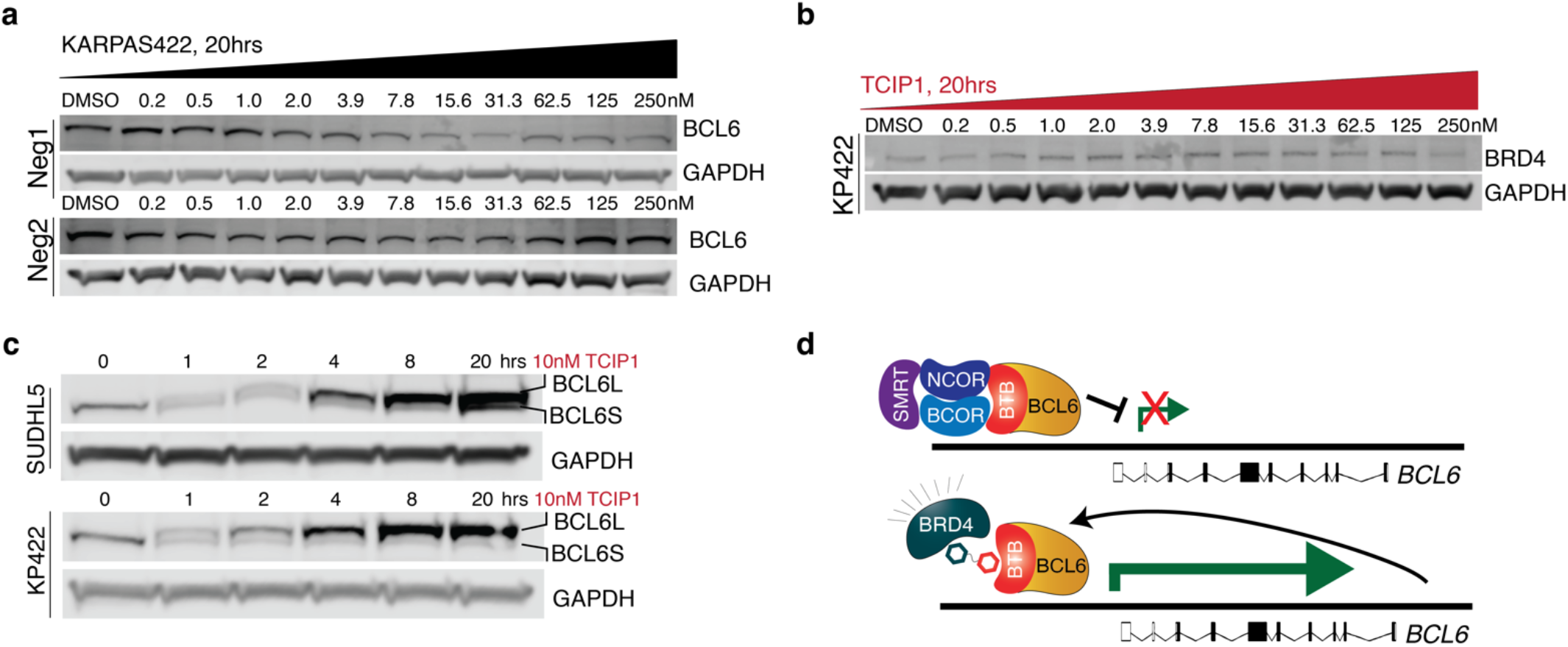
Conversion of BCL6 auto-inhibitory pathway to a feedforward loop. **a.** Control Neg1 and Neg2 effect on BCL6 protein levels at 20hrs treatment in KARPAS422 cells. **b.** Effect on BRD4 levels at 20hrs treatment with TCIP1 in KARPAS422 cells. **c.** Kinetics of BCL6 upregulation in two separate DLBCL cell lines, KARPAS422 and SUDHL5, after addition of 10 nM TCIP1. **d.** Model for conversion of BCL6 auto-inhibitory circuit to a positive feedback loop.

**Extended Data Figure 7.**
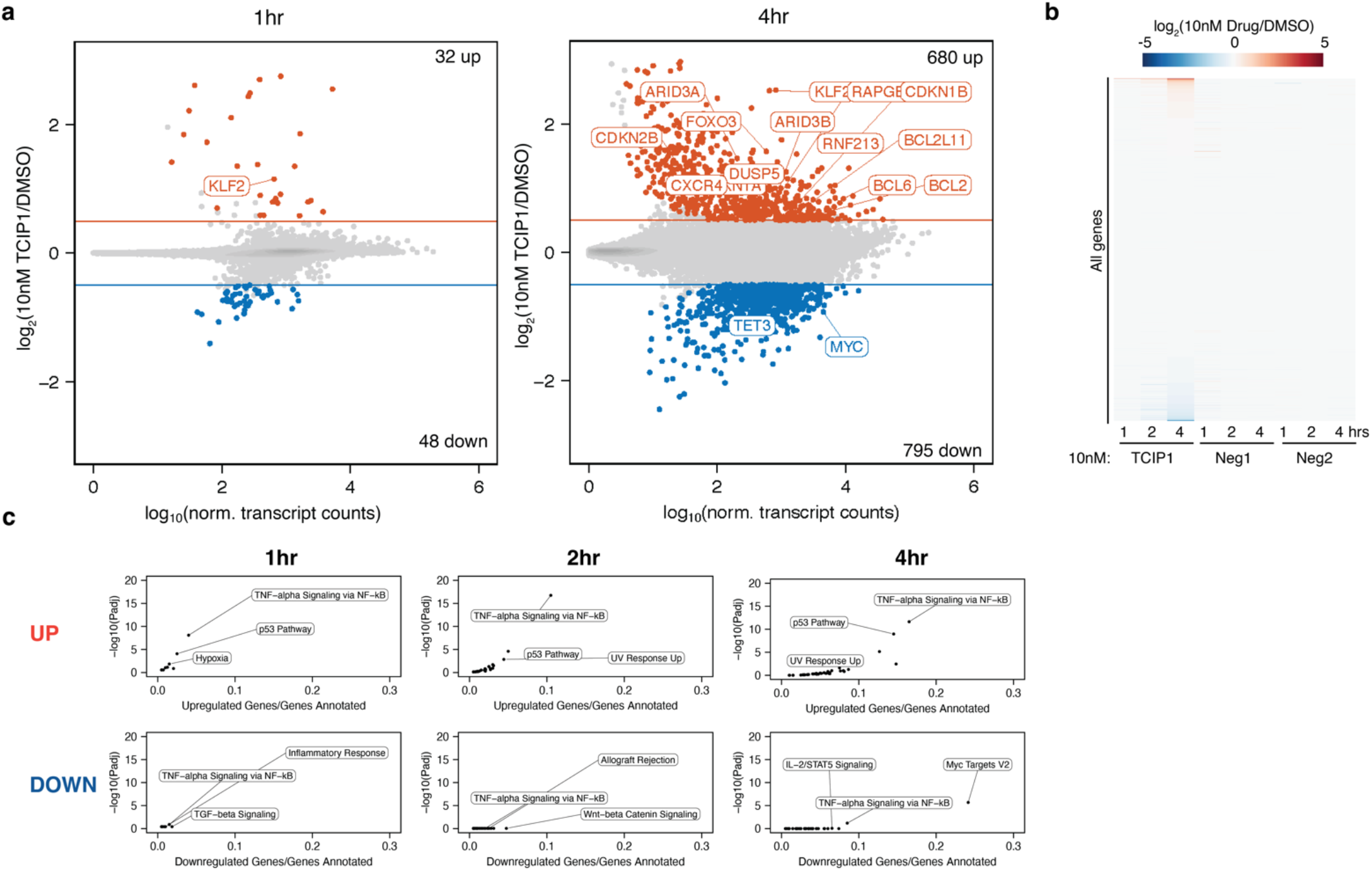
Specific activation of gene expression by TCIP1 but not related controls. **a.** Gene expression changes after 1hr or 4hrs addition of 10 nM TCIP1 in KARPAS422 cells. Significance cutoffs were p_adj_≤0.05 and |log_2_(Drug/DMSO)|≥0.5), n=3 biological repeats. **b.** Specific effects of TCIP1 across transcriptome. For Neg1 and Neg2, n=2 biological repeats. **c.** Enrichment analysis of upregulated and downregulated genes.

**Extended Data Figure 8.**
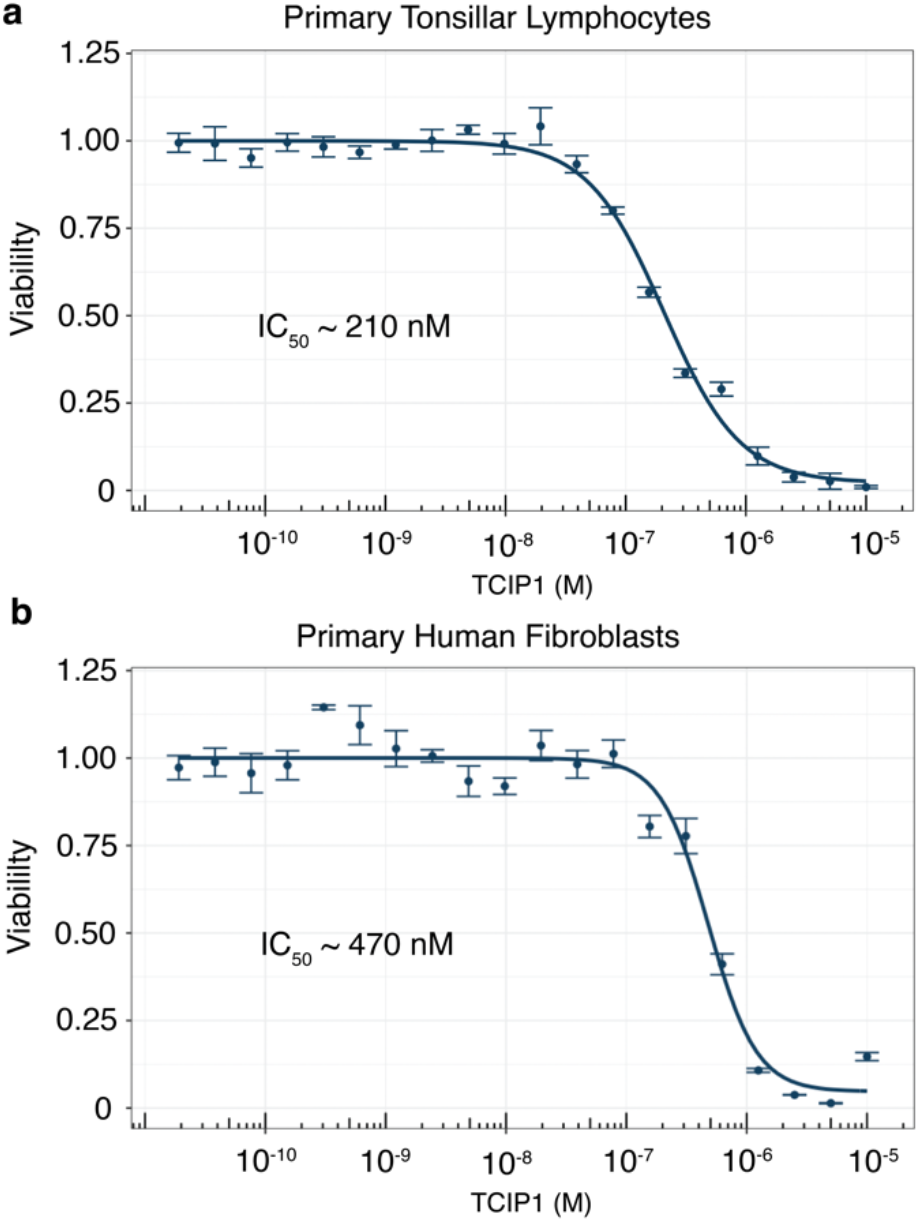
TCIP1 toxicity studies in primary human cells. **a.** Effect of TCIP1 on cell viability of primary human tonsillar lymphocytes. **b.** Effect of TCIP1 on cell viability of primary fibroblasts. n=3 biological repeats, mean±s.d.

**Extended Data Figure 9.**
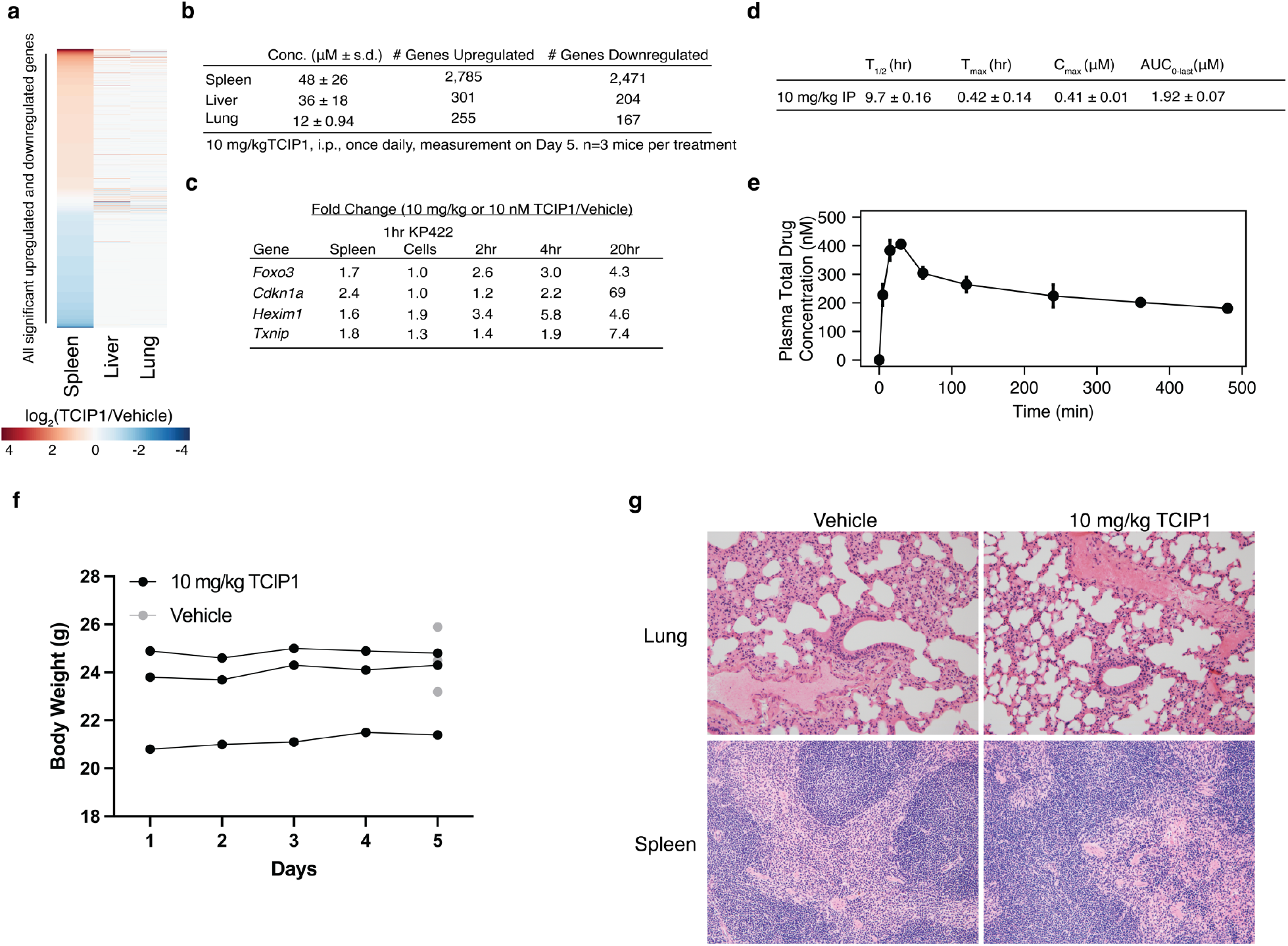
TCIP1 pharmacokinetic properties, toxicity, and context-specific target engagement in mice. **a.** Tissue-specific transcriptomic effects of TCIP1, treated at 10 mg/kg IP q.d. for 5 days. n=3 mice each for treatment and vehicle. **b.** Quantification of transcriptome changes in the liver, lung, and spleen and associated accumulated tissue concentrations of TCIP1. **c**. Comparison of key gene targets upregulated by TCIP1 in both cultured DLBCL cells (KARPAS422 a.k.a KP422) and in the Spleen. **d**. Pharmacokinetic parameters of TCIP1. **e**. Plasma concentrations of total TCIP1 over time after treatment. **f.** Body weight of treated mice. No adverse effects were noticed. **g**. H&E staining of lung and spleen from representative vehicle and drug-treated mice.

