## Supplemental Information for "Rewiring Cancer Drivers to Activate Apoptosis"

**A. Supplemental Methods**
  1. **Chemical Synthesis**
  2. **Protein Constructs and Purification**

**B. Supplemental Figures**
  1. **Uncropped blots**
  2. **Coomassie-stained Gels**
  3. **Flow cytometry gating strategy**

#### Chemical Synthesis

Unless otherwise noted, reagents and solvents were obtained from commercial suppliers and were used without further purification. $^{1}$H NMR spectra were recorded on 500 MHz (Bruker A500), and chemical shifts are reported in parts per million (ppm, d) downfield from tetramethylsilane (TMS). Coupling constants (J) are reported in Hz. Spin multiplicities are described as s (singlet), br (broad singlet), d (doublet), t (triplet), q (quartet), and m (multiplet). Mass spectra were obtained on a Waters Micromass ZQ instrument. Preparative HPLC was performed on a Waters Sunfire C18 column (19 x 50 mm, 5μM) using a gradient of 15-95% methanol in water containing 0.05% trifluoroacetic acid (TFA) over 22 min (28 min run time) at a flow rate of 20 mL/min. Purities of assayed compounds were in all cases greater than 95%, as determined by reverse-phase HPLC analysis.

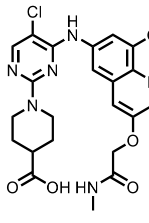

S1

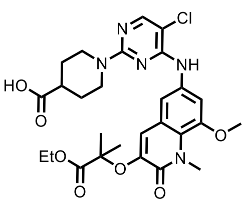

S2

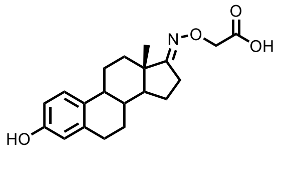

S3

S1 were prepared according to the known literature procedures. [1]
S2 were prepared according to the known literature procedures [2]
S3 were prepared according to the known literature procedures. [3]

**Synthesis of (S)-1-(5-chloro-4-((8-methoxy-1-methyl-3-(2-(methylamino)-2-oxoethoxy)-2-oxo-1,2-dihydroquinolin-6-yl)amino)pyrimidin-2-yl)-N-(7-(2-(4-(4-chlorophenyl)-2,3,9-trimethyl-6H-thieno[3,2-f][1,2,4]triazolo[4,3-a][1,4]diazepin-6-yl)acetamido)heptyl)piperidine-4-carboxamide (TCIP 1).**

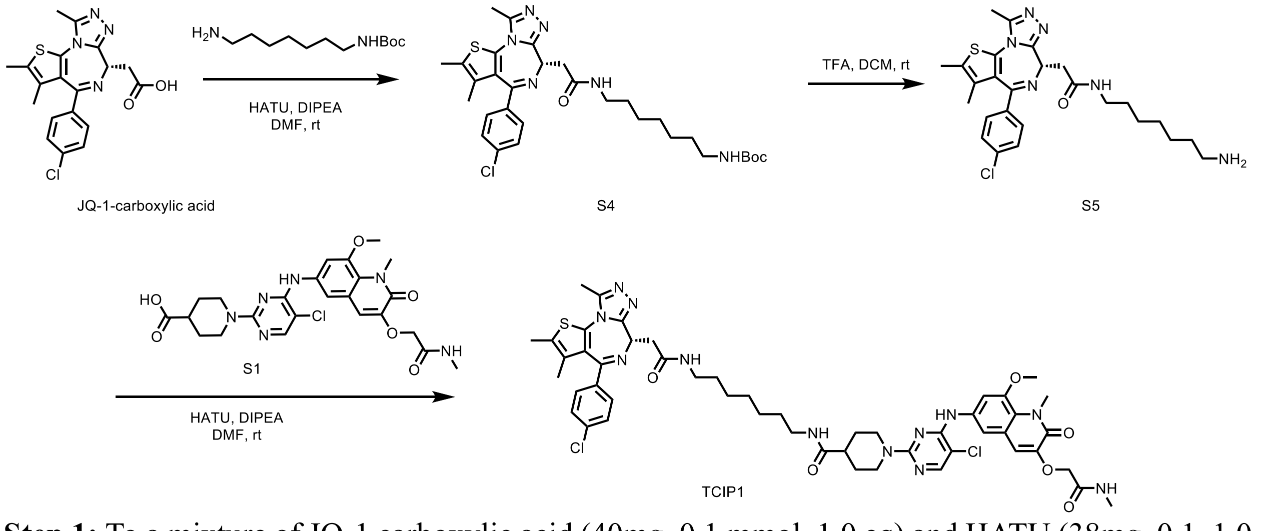


**Step 1:** To a mixture of JQ-1 carboxylic acid (40mg, 0.1 mmol, 1.0 eq) and HATU (38mg, 0.1, 1.0 eq) DIPEA (52ul, 0.3 mmol, 3eq) in DMF( 2mL) was added tert-butyl (7-aminoheptyl)carbamate ( 27mg, 0.12 mmol, 1.2eq), the mixture was stirred at room temperature for 1 hour. LC-MS indicated formation of desired product. The mixture was concentrated and purified via prep-HPLC to afford intermediate **S4** (51mg). LC-MS (ESI) m/z : [M+H]+: calcd 613.27 , found 613.38.

**Step 2:** To intermediate **S4**(51mg) was added a solution of DCM/ Trifluoroacetic acid ( 1 mL:0.3 mL). The mixture was stirred at room temperature for 1 hour. LC-MS indicated formation of desired product. The mixture was concentrated under reduced pressure to give crude product **S5** which was used directly without future purification. LC-MS (ESI) m/z : [M+H]+: calcd 513.22 , found 513.41.

**Step 3:**To a mixture of **S5** (10mg, 0.02mmol, 1.0 eq), HATU (7.6mg, 0.02 mmol, 1.0 eq), DIPEA (10uL, 0.06mmol, 3eq) in DMF was added intermediate **S1**(10.6mg, 0.06 mmol, 1 eq), the mixture was stirred at room temperature for 1 hour. LC-MS indicated formation of desired product. The mixture was concentrated and purified via prep-HPLC to afford desired product **TCIP1** (11mg) LC-MS (ESI) m/z : [M+H]+: calcd 1025.38 , found 1025.40.

**$^1$H NMR** (500 MHz, DMSO-d6) δ 8.96 (s, 1H), 8.09 (t, J = 5.7 Hz, 1H), 8.03 (s, 1H), 7.88 (d, J = 5.2 Hz, 1H), 7.70 (t, J = 5.6 Hz, 1H), 7.44 (s, 2H), 7.41 (d, J = 8.2 Hz, 2H), 7.34 (d, J = 8.2 Hz, 2H), 6.94 (s, 1H), 4.48 (s, 2H), 4.46 – 4.35 (m, 3H), 3.79 (s, 3H), 3.77 (s, 3H), 3.14 (qd, J = 15.0, 7.1 Hz, 2H), 3.02 (dq, J = 13.0, 6.7 Hz, 2H), 2.94 (q, J = 6.7 Hz, 2H), 2.84 (t, J = 12.5 Hz, 2H), 2.58 (d, J = 4.6 Hz, 3H), 2.52 (s, 3H), 2.33 (s, 3H), 1.64 (d, J = 12.8 Hz, 2H), 1.55 (s, 3H), 1.49 – 1.39 (m, 2H), 1.36 – 1.30 (m, 3H), 1.25 – 1.11 (m, 6H).

**Synthesis of (R)-1-(5-chloro-4-((8-methoxy-1-methyl-3-(2-(methylamino)-2-oxoethoxy)-2-oxo-1,2-dihydroquinolin-6-yl)amino)pyrimidin-2-yl)-N-(7-(2-(4-(4-chlorophenyl)-2,3,9-trimethyl-6H-thieno[3,2-f][1,2,4]triazolo[4,3-a][1,4]diazepin-6-yl)acetamido)heptyl)piperidine-4-carboxamide (Neg1).**

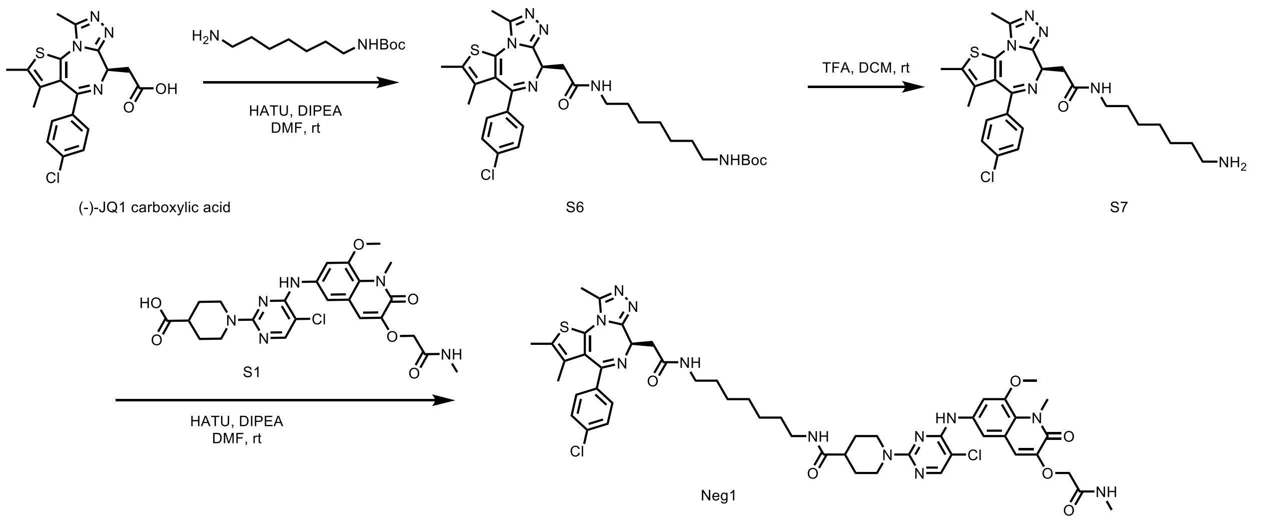


**Step 1:** To a mixture of (-)-JQ1 carboxylic acid (20mg, 0.05 mmol, 1.0 eq) and HATU (20mg, 0.05, 1.0 eq) DIPEA (26ul, 0.15 mmol, 3eq) in DMF( 1mL) was added tert-butyl (7-aminoheptyl)carbamate ( 13mg, 0.06 mmol, 1.2eq), the mixture was stirred at room temperature for 1 hour. LC-MS indicated formation of desired product. The mixture was concentrated and purified via prep-HPLC to afford intermediate **S6** (22mg) . LC-MS (ESI) m/z : [M+H]+: calcd 613.27 , found 613.29.

**Step 2:** To intermediate **S6**(22mg) was added a solution of DCM/ Trifluoroacetic acid ( 1 mL:0.3 mL). The mixture was stirred at room temperature for 1 hour. LC-MS indicated formation of desired product. The mixture was concentrated under reduced pressure to give crude product **S7**

which was used directly without future purification. LC-MS (ESI) m/z : [M+H]+: calcd 513.22 , found 513.36.

**Step 3:** To a mixture of **S7** (10mg, 0.02mmol, 1.0 eq), HATU (7.6mg, 0.02 mmol, 1.0 eq), DIPEA (10uL, 0.06mmol, 3eq) in DMF was added intermediate **S1** (10.6mg, 0.06 mmol, 1.0 eq), the mixture was stirred at room temperature for 1 hour. LC-MS indicated formation of desired product. The mixture was concentrated and purified via prep-HPLC to afford desired product **Neg1** (9mg)

LC-MS (ESI) m/z : [M+H]+: calcd 1025.38 , found 1025.42.

**$^1$H NMR** (500 MHz, DMSO-d6) δ 9.12 (s, 1H), 8.16 (t, J = 5.7 Hz, 1H), 8.11 (s, 1H), 8.02 (t, J = 4.8 Hz, 1H), 7.78 (t, J = 5.6 Hz, 2H), 7.51 (s, 2H), 7.48 (d, J = 8.6 Hz, 2H), 7.44 – 7.40 (m, 1H), 7.03 (s, 1H), 4.55 (s, 2H), 4.50 – 4.40 (m, 3H), 3.87 (s, 3H), 3.86 (s, 3H), 3.28 – 2.88 (m, 8H), 2.65 (d, J = 4.6 Hz, 3H), 2.60 (s, 3H), 2.40 (s, 3H), 1.71 (dd, J = 13.5, 3.7 Hz, 2H), 1.62 (s, 3H), 1.56 –1.46 (m, 2H), 1.45 – 1.33 (m, 3H), 1.30 – 1.19 (m, 6H).

**Synthesis of (S)-2-((6-((5-chloro-2-(4-((7-(2-(4-(4-chlorophenyl)-2,3,9-trimethyl-6H-thieno[3,2-f][1,2,4]triazolo[4,3-a][1,4]diazepin-6-yl)acetamido)heptyl)carbamoyl)piperidin-1-yl)pyrimidin-4-yl)amino)-8-methoxy-1-methyl-2-oxo-1,2-dihydroquinolin-3-yl)oxy)-2-methylpropanoic acid (Neg2).**

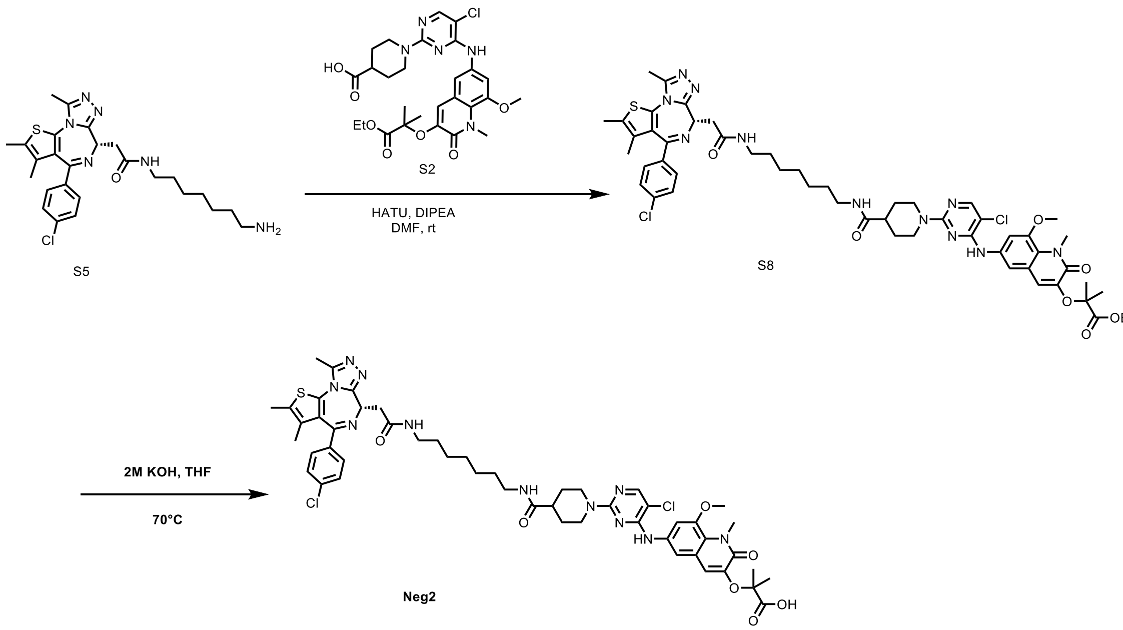


**Step 1:** To a mixture of **S5** (15mg, 0.03mmol, 1.0 eq), HATU (11.4mg, 0.03 mmol, 1.0 eq), DIPEA (15uL, 0.09mmol, 3eq) in DMF was added intermediate **S2** (17.1mg, 0.03 mmol, 1.0 eq), the mixture was stirred at room temperature for 1 hour. LC-MS indicated formation of desired product. The mixture was concentrated and purified via prep-HPLC to afford desired product **S9** (16 mg)

LC-MS (ESI) m/z : [M+H]+: calcd 1068.41 , found 1068.53.

**Step 2:** To a mixture of S9 (16mg) in THF (0.5 mL) was added 2M KOH (0.5 mL). The mixture was stirred at 70°C for 6 h. The resulting mixture was cooled to room temperature, acidified to PH=5~7 with 2M HCl, exacted with EtOAc and the organic layer was washed with water and brine. The organic layer was removed and purified by HPLC to give **Neg2** (10.1 mg) as white solid. LC-MS (ESI) m/z : [M+H]+: calcd 1040.38 , found 10.40.53

**$^{1}$H NMR** (500 MHz, DMSO-d6) δ 9.12 (s, 1H), 8.16 (t, J = 5.7 Hz, 1H), 8.12 (s, 1H), 7.78 (t, J = 5.6 Hz, 1H), 7.59 (d, J = 2.3 Hz, 1H), 7.53 (d, J = 2.2 Hz, 1H), 7.48 (d, J = 8.8 Hz, 2H), 7.44 – 7.40 (m, 2H), 6.91 (s, 1H), 4.53 – 4.47 (m, 2H), 4.49 – 4.42 (m, 3H), 3.87 (s, 3H), 3.85 (s, 3H), 3.28 – 2.89 (m, 8H), 2.60 (s, 3H), 2.41 (s, 3H), 1.77 – 1.69 (m, 2H), 1.62 (s, 3H), 1.55 (s, 6H), 1.51 (d, J = 3.9 Hz, 1H), 1.45-1.39 (m, 2H), 1.40 – 1.35 (m, 2H), 1.30-1.18 (m, 6H).

**Synthesis of 1-(5-chloro-4-((8-methoxy-1-methyl-3-(2-(methylamino)-2-oxoethoxy)-2-oxo-1,2-dihydroquinolin-6-yl)amino)pyrimidin-2-yl)-N-(2-(2-(2-(2-((((13S,E)-3-hydroxy-13-methyl-6,7,8,9,11,12,13,14,15,16-decahydro-17H-cyclopenta[a]phenanthren-17-ylidene)amino)oxy)acetamido)ethoxy)ethoxy)ethyl)piperidine-4-carboxamide (TCIP2).**

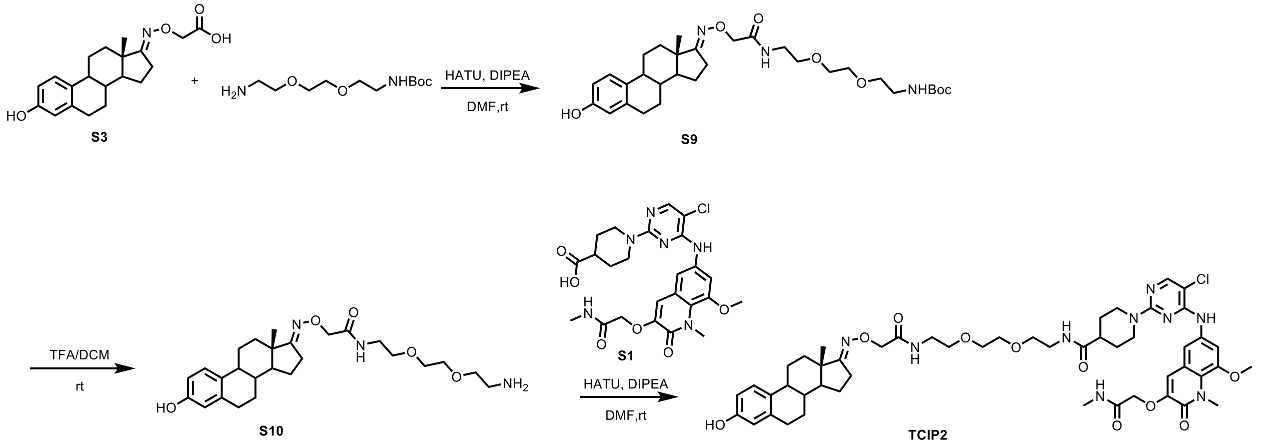


**Step 1**: A solution S3(20 mg, 0.06 mmol) t-Boc-N-amido-PEG2-amine and (15 mg, 0.06 mmol), HATU (33 mg, 0.09 mmol) and DIPEA (33 uL, 0.2 mmol) in DMF (0.5 mL) was stirred at room temperature for 1 h. The solvent was removed under reduced pressure and was purified by flash chromatography to afford coupling product **S9** (24 mg) as white solid. LC-MS (ESI) m/z : [M+H]+: calcd 574.35 , found 574.41.

**Step 2**: **S9** (24 mg, 0.04 mmol) was dissolved in 1 mL DCM and subjected to 0.2 mL TFA at room temperature for 1 h. The solvent was removed under reduced pressure and the crude **S10** was used in the next step without further purification. LC-MS (ESI) m/z : [M+H]+: calcd 474.30 , found 474.34.

**Step 3**: To a mixture of **S10** (10 mg,0.02mmol), **S1** (13 mg, 0.024 mmol) and DIPEA (14 uL, 0.08 mmol) in DMF (1 mL)was added HATU (9 mg, 0.024mmol). The resultant mixture was stirred at room temperature for 2h. The solvent was removed under reduced pressure and purified by HPLC to give **TCIP2** (10 mg) as white solid. LC-MS (ESI) m/z : [M+H]+: calcd 986.45 , found 986.42. 1H NMR (500 MHz, DMSO) δ 8.93 (s, 1H), 8.80 (s, 1H), 8.00 (s,1H), 7.89 (d, J = 5.3 Hz, 2H), 7.78 (t, J = 5.6 Hz, 2H), 7.46 (s, 2H), 7.34 (q, J = 9.6 Hz, 2H),7.00 - 6.91 (m, 3H), 6.42 (d, J= 8.3 Hz, 2H), 6.37 (d, J =1.1 Hz, 2H), 4.45 (d, J = 29.5 Hz,5H), 4.26 (s, 2H), 3.79 (d, J = 6.7 Hz, 7H), 3.33 (dt, J = 16.2, 5.8 Hz, 11H), 3.19 (q, J = 5.9 Hz, 5H), 3.11 (q, J= 6.0 Hz, 4H), 2.83 (t, J= 12.4 Hz, 4H), 2.73-2.61 (m, 4H), 2.58 (d, J = 4.7 Hz, 4H), 2.48 (d, J = 9.6 Hz, 4H), 2.38 – 2.29 (m, 3H), 2.20 (d, J= 13.2 Hz, 2H), 2.06 (t, J= 11.8 Hz, 2H), 1.82 –1.73 (m, 4H), 1.63 – 1.24 (m, 15H), 1.24 –1.15 (m, 4H), 0.79 (s, 3H).

**General synthesis 1:**

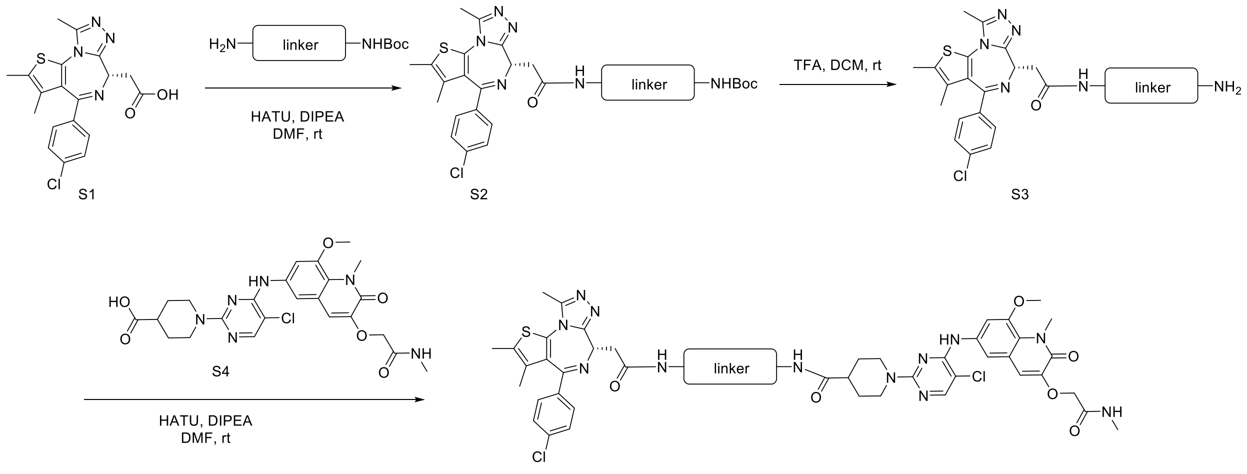


**Step 1: Synthesis of intermediate S2.**
To a mixture of JQ-1 carboxylic acid 1 (1.0 eq) and HATU (1.0 eq) DIPEA (3eq) in DMF was added linker( 1.2eq), the mixture was stirred at room temperature for 1 hour. LC-MS indicated formation of desired product. The mixture was concentrated and purified via prep-HPLC to afford intermediate **S2**.

**Step 2: Synthesis of intermediate S3.**
To intermediate **S2** was added a solution of DCM/ Trifluoroacetic acid ( 3:1). The mixture was stirred at room temperature for 1 hour. LC-MS indicated formation of desired product. The mixture was concentrated under reduced pressure to give crude product which was used directly without future purification.

**Step 3:Synthesis of desired product.**
To a mixture of 1-(5-chloro-4-((8-methoxy-1-methyl-3-(2-(methylamino)-2-oxoethoxy)-2-oxo-1,2-dihydroquinolin-6-yl)amino)pyrimidin-2-yl)piperidine-4-carboxylic acid (**S4**) (1.0 eq), HATU (1.0 eq), DIPEA (3eq) in DMF was added intermediate **S3** ( 1.2 eq), the mixture was stirred at room temperature for 1 hour. LC-MS indicated formation of desired product. The mixture was concentrated and purified via prep-HPLC to afford desired product.

**General synthesis 2:**

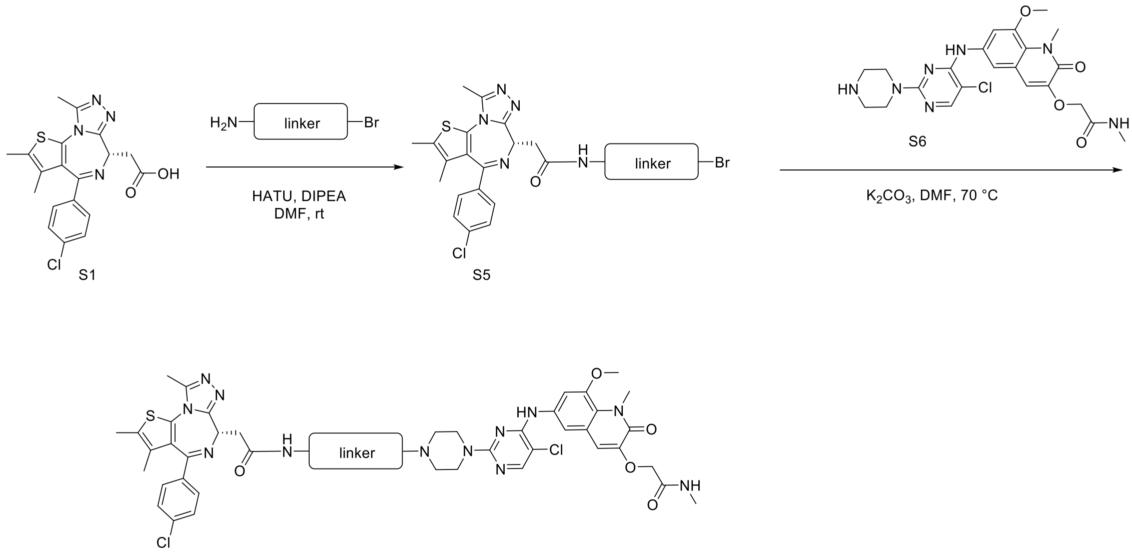


**Step 1: Synthesis of intermediate S5.**

To a mixture of JQ-1 carboxylic acid 1 (1.0 eq) and HATU (1.0 eq) DIPEA (3eq) in DMF was added linker( 1.2eq), the mixture was stirred at room temperature for 1 hour. LC-MS indicated formation of desired product. The mixture was concentrated and purified via prep-HPLC to afford intermediate **S5**.

**Step 2: Synthesis of desired product.**

To a mixture of 2-((6-((5-chloro-2-(piperazin-1-yl)pyrimidin-4-yl)amino)-8-methoxy-1-methyl-2-oxo-1,2-dihydroquinolin-3-yl)oxy)-N-methylacetamide (**S6**) (1.0 eq), $K_2CO_3$ (3.0 eq) in DMF was added **intermediate S5** ( 1.3 eq), the mixture was stirred at 70 °C for 1 hour. LC-MS indicated formation of desired product. The mixture was concentrated and purified via prep-HPLC to afford desired product.

**Characterization of compounds.**

(S)-1-(5-chloro-4-((8-methoxy-1-methyl-3-(2-(methylamino)-2-oxoethoxy)-2-oxo-1,2-dihydroquinolin-6-yl)amino)pyrimidin-2-yl)-N-(6-(2-(4-(4-chlorophenyl)-2,3,9-trimethyl-6H-thieno[3,2-f][1,2,4]triazolo[4,3-a][1,4]diazepin-6-yl)acetamido)hexyl)piperidine-4-carboxamide(**JWZ-7-6**)

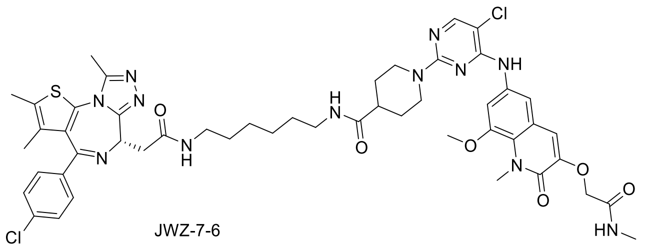

JWZ-7-6

**LC-MS** (ESI) m/z: 1011.4 $[M+H]^+$.

**1H NMR** (500 MHz, DMSO-d6) δ 8.98 (s, 1H), 8.09 (t, J = 5.8 Hz, 1H), 8.03 (d, J = 1.7 Hz, 1H), 7.88 (q, J = 4.8 Hz, 1H), 7.71 (t, J = 5.7 Hz, 1H), 7.44 (d, J = 1.8 Hz, 2H), 7.40 (d, J = 7.9 Hz, 2H), 7.36 – 7.31 (m, 2H), 6.94 (s, 1H), 4.48 (s, 2H), 4.43 – 4.37 (m, 3H), 3.79 (s, 3H), 3.78 (s, 3H), 3.17 (dd, J = 15.0, 8.3 Hz, 1H), 3.13 – 3.03 (m, 2H), 3.02 – 2.97 (m, 1H), 2.94 (q, J = 6.6 Hz, 2H), 2.85 (t, J = 12.6 Hz, 2H), 2.58 (d, J = 4.6 Hz, 3H), 2.52 (d, J = 1.7 Hz, 3H), 2.32 (s, 3H), 1.64 (d, J = 12.9 Hz, 2H), 1.53 (s, 3H), 1.48 – 1.27 (m, 6H), 1.24 – 1.16 (m, 4H).

(S)-1-(5-chloro-4-((8-methoxy-1-methyl-3-(2-(methylamino)-2-oxoethoxy)-2-oxo-1,2-dihydroquinolin-6-yl)amino)pyrimidin-2-yl)-N-(2-(4-(2-(2-(4-(4-chlorophenyl)-2,3,9-trimethyl-6H-thieno[3,2-f][1,2,4]triazolo[4,3-a][1,4]diazepin-6-yl)acetamido)ethyl)piperazin-1-yl)ethyl)piperidine-4-carboxamide( **JWZ-7-13**)

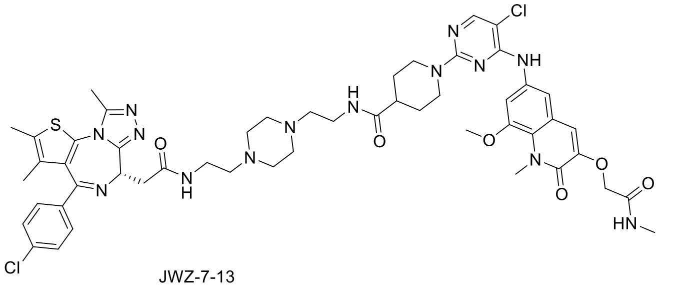


JWZ-7-13

**LC-MS** (ESI) m/z: 1067.4 $[M+H]^+$.

**1H NMR** (500 MHz, DMSO-d6) δ 8.90 (s, 1H), 8.34 (s, 1H), 8.02 (s, 1H), 7.96 (s, 1H), 7.90 (q, J = 4.4 Hz, 1H), 7.49 – 7.44 (m, 2H), 7.42 (d, J = 8.3 Hz, 2H), 7.35 (d, J = 8.2 Hz, 2H), 6.94 (s, 1H), 4.48 (s, 2H), 4.46 – 4.40 (m, 6H), 3.80 (s, 3H), 3.79 (s, 3H), 3.43 – 3.15 (m, 10H), 2.97 – 2.79 (m, 8H), 2.58 (d, J = 4.6 Hz, 3H), 2.53 (s, 3H), 2.34 (s, 3H), 1.68 (d, J = 12.6 Hz, 2H), 1.55 (s, 3H), 1.43 (q, J = 12.1, 11.6 Hz, 2H).

(S)-1-(5-chloro-4-((8-methoxy-1-methyl-3-(2-(methylamino)-2-oxoethoxy)-2-oxo-1,2-dihydroquinolin-6-yl)amino)pyrimidin-2-yl)-N-(3-(4-(3-(2-(4-(4-chlorophenyl)-2,3,9-trimethyl-6H-thieno[3,2-f][1,2,4]triazolo[4,3-a][1,4]diazepin-6-yl)acetamido)propyl)piperazin-1-yl)propyl)piperidine-4-carboxamide( **JWZ-7-14**)

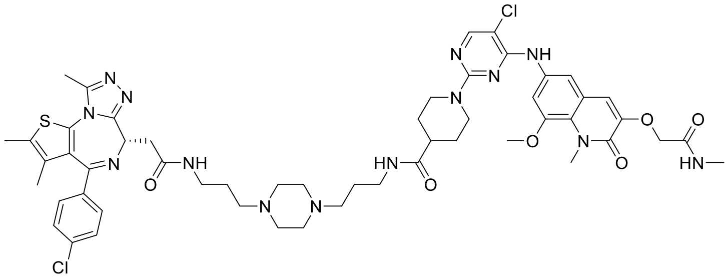


JWZ-7-14

**LC-MS** (ESI) m/z: 1067.4 $[M+H]^+$.

**1H NMR** (500 MHz, DMSO-d6) δ 8.83 (s, 1H), 8.27 (d, J = 6.4 Hz, 1H), 8.02 (d, J = 2.9 Hz, 1H), 7.90 (d, J = 5.5 Hz, 2H), 7.49 (d, J = 2.4 Hz, 1H), 7.45 (d, J = 2.5 Hz, 1H), 7.43 (dd, J = 8.6, 2.9 Hz, 2H), 7.38 – 7.34 (m, 2H), 6.93 (d, J = 3.0 Hz, 1H), 4.49 (d, J = 2.8 Hz, 2H), 4.47 – 4.41 (m, 3H), 3.80 (s, 3H), 3.79 (s, 3H), 3.27 – 3.12 (m, 5H), 3.04 (q, J = 6.7, 6.1 Hz, 4H), 2.99 – 2.79 (m, 8H), 2.58 (d, J = 4.5 Hz, 3H), 2.53 (d, J = 2.9 Hz, 3H), 2.35 (d, J = 2.9 Hz, 3H), 2.30 (s, 1H), 1.75 – 1.61 (m, 6H), 1.56 (d, J = 2.8 Hz, 3H), 1.50 – 1.37 (m, 3H), 1.23 – 1.13 (m, 2H).

(S)-1-(5-chloro-4-((8-methoxy-1-methyl-3-(2-(methylamino)-2-oxoethoxy)-2-oxo-1,2-dihydroquinolin-6-yl)amino)pyrimidin-2-yl)-N-(9-(2-(4-(4-chlorophenyl)-2,3,9-trimethyl-6H-thieno[3,2-f][1,2,4]triazolo[4,3-a][1,4]diazepin-6-yl)acetamido)nonyl)piperidine-4-carboxamide(**JWZ-7-15**)

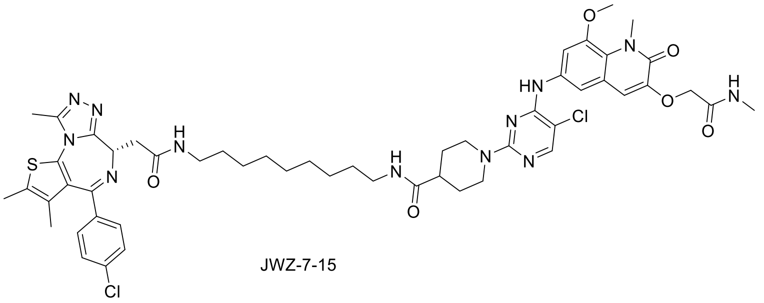

JWZ-7-15

**LC-MS** (ESI) m/z: 1053.4 $[M+H]^+$.

**1H NMR** (500 MHz, DMSO-d6) δ 8.86 (s, 1H), 8.08 (t, J = 5.7 Hz, 1H), 8.01 (s, 1H), 7.89 (d, J = 4.7 Hz, 1H), 7.69 (t, J = 5.6 Hz, 1H), 7.46 (s, 2H), 7.40 (d, J = 8.8 Hz, 2H), 7.37 – 7.32 (m, 2H), 6.93 (s, 1H), 4.48 (s, 2H), 4.45 – 4.39 (m, 3H), 3.80 (s, 3H), 3.78 (s, 3H), 3.22 – 2.97 (m, 7H), 2.93 (q, J = 6.6 Hz, 3H), 2.83 (td, J = 12.9, 2.7 Hz, 3H), 2.58 (d, J = 4.7 Hz, 3H), 2.52 (s, 3H), 2.33 (s, 3H), 2.30 (dt, J = 7.5, 3.7 Hz, 1H), 1.66 – 1.60 (m, 2H), 1.55 (s, 3H), 1.47 – 1.26 (m, 8H), 1.22 –1.10 (m, 8H).

(S)-1-(5-chloro-4-((8-methoxy-1-methyl-3-(2-(methylamino)-2-oxoethoxy)-2-oxo-1,2-dihydroquinolin-6-yl)amino)pyrimidin-2-yl)-N-(10-(2-(4-(4-chlorophenyl)-2,3,9-trimethyl-6H-thieno[3,2-f][1,2,4]triazolo[4,3-a][1,4]diazepin-6-yl)acetamido)decyl)piperidine-4-carboxamide (**JWZ-7-20**)

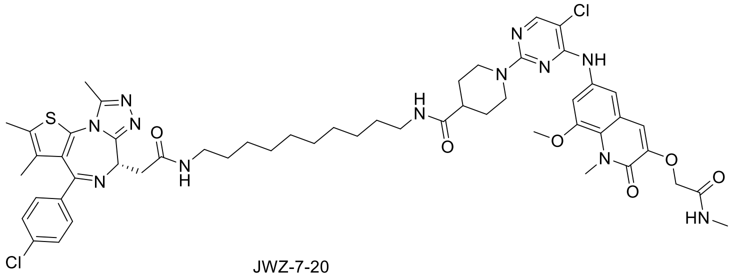

JWZ-7-20

**LC-MS** (ESI) m/z: 1053.4 $[M+H]^+$.

**1H NMR** (500 MHz, DMSO-d6) δ 9.02 (s, 1H), 8.16 (t, J = 5.7 Hz, 1H), 8.10 (s, 1H), 7.97 (d, J = 4.7 Hz, 1H), 7.76 (t, J = 5.6 Hz, 1H), 7.53 (s, 2H), 7.50 – 7.41 (m, 4H), 7.01 (s, 1H), 4.55 (s, 2H), 4.53 – 4.45 (m, 3H), 3.87 (s, 3H), 3.86 (s, 3H), 3.25 (dd, J = 14.9, 8.5 Hz, 2H), 3.20 – 3.09 (m, 3H), 3.09 – 2.98 (m, 4H), 2.92 (td, J = 13.0, 2.8 Hz, 3H), 2.65 (d, J = 4.6 Hz, 3H), 2.60 (s, 3H), 2.41 (s, 3H), 2.39 – 2.35 (m, 1H), 1.74 – 1.68 (m, 2H), 1.62 (s, 3H), 1.51 (dd, J = 12.1, 3.8 Hz, 3H), 1.43 (t, J = 7.1 Hz, 2H), 1.36 (t, J = 6.8 Hz, 2H), 1.29 – 1.16(m, 11H).

(S)-2-((6-((5-chloro-2-(4-(11-(2-(4-(4-chlorophenyl)-2,3,9-trimethyl-6H-thieno[3,2-f][1,2,4]triazolo[4,3-a][1,4]diazepin-6-yl)acetamido)undecyl)piperazin-1-yl)pyrimidin-4-yl)amino)-8-methoxy-1-methyl-2-oxo-1,2-dihydroquinolin-3-yl)oxy)-N-methylacetamide (**JWZ-7-22**)

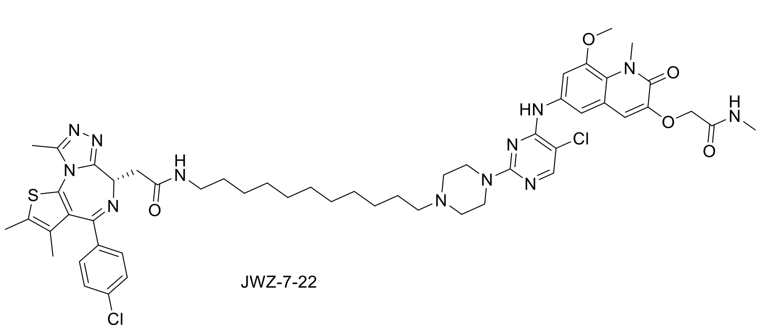


JWZ-7-22

**LC-MS** (ESI) m/z: 1039.4 $[M+H]^+$.

**1H NMR** (500 MHz, DMSO-d6) δ 9.66 (s, 1H), 9.01 (s, 1H), 8.16 (d, J = 3.2 Hz, 1H), 8.00 (q, J = 4.7 Hz, 1H), 7.56 (d, J = 2.3 Hz, 1H), 7.48 (d, J = 8.7 Hz, 2H), 7.46 – 7.41 (m, 2H), 7.08 (s, 1H), 4.57 (s, 2H), 4.50 (dd, J = 8.3, 5.8 Hz, 3H), 3.88 (s, 3H), 3.87 (s, 3H), 3.32 – 2.99 (m, 12H), 2.66 (d, J = 4.6 Hz, 3H), 2.60 (s, 3H), 2.41 (s, 3H), 1.69 – 1.64 (m, 2H)1.63 (s, 3H), 1.44 (t, J = 7.0 Hz, 2H), 1.35 – 1.20 (m, 14H).

(S)-2-((6-((5-chloro-2-(4-(8-(2-(4-(4-chlorophenyl)-2,3,9-trimethyl-6H-thieno[3,2-f][1,2,4]triazolo[4,3-a][1,4]diazepin-6-yl)acetamido)octyl)piperazin-1-yl)pyrimidin-4-yl)amino)-8-methoxy-1-methyl-2-oxo-1,2-dihydroquinolin-3-yl)oxy)-N-methylacetamide( **JWZ-7-23**)

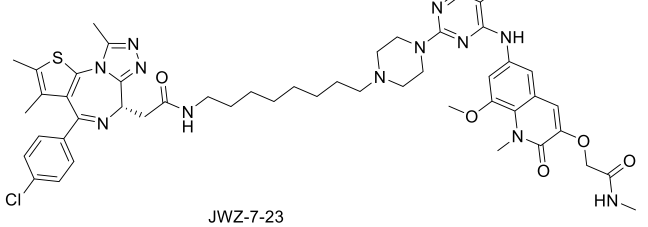


JWZ-7-23

**LC-MS** (ESI) m/z: 997.4 $[M+H]^+$.

**1H NMR** (500 MHz, DMSO-d6) δ 9.65 (s, 1H), 8.93 (s, 1H), 8.10 (d, J = 5.7 Hz, 1H), 8.08 (s, 1H), 7.91 (q, J = 4.6 Hz, 1H), 7.48 (d, J = 2.3 Hz, 1H), 7.42 (d, J = 1.6 Hz, 1H), 7.40 (s, 1H), 7.38 – 7.33 (m, 2H), 7.00 (s, 1H), 4.50 (s, 2H), 4.47 – 4.40 (m, 3H), 3.80 (s, 3H), 3.80 (s, 3H), 3.22 – 3.10 (m, 6H), 3.06 – 2.92 (m, 6H), 2.58 (d, J = 4.7 Hz, 3H), 2.52 (s, 3H), 2.34 (s, 3H), 1.60 (d, J = 9.2 Hz, 2H), 1.55 (s, 3H), 1.36 (d, J = 6.4 Hz, 2H), 1.22 (s, 6H), 1.17 (s, 2H).

(S)-2-((6-((5-chloro-2-(4-(1-(4-(4-chlorophenyl)-2,3,9-trimethyl-6H-thieno[3,2-f][1,2,4]triazolo[4,3-a][1,4]diazepin-6-yl)-2-oxo-6,9,12-trioxa-3-azatetradecan-14-yl)piperazin-1-yl)pyrimidin-4-yl)amino)-8-methoxy-1-methyl-2-oxo-1,2-dihydroquinolin-3-yl)oxy)-N-methylacetamide (**JWZ-7-24**)

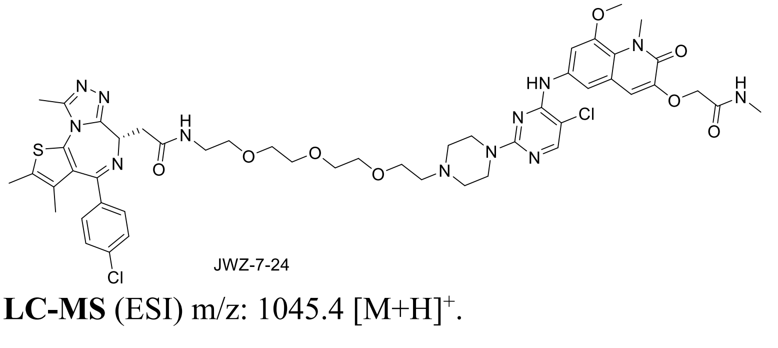


JWZ-7-24

**LC-MS** (ESI) m/z: 1045.4 $[M+H]^+$.

**1H NMR** (500 MHz, DMSO-d6) δ 9.85 (s, 1H), 9.00 (s, 1H), 8.25 (t, J = 5.7 Hz, 1H), 8.14 (s, 1H), 8.04 (q, J = 4.7 Hz, 1H), 7.53 (d, J = 2.3 Hz, 1H), 7.48 (d, J = 8.8 Hz, 2H), 7.45 – 7.39 (m, 2H), 7.08 (s, 1H), 4.56 (s, 2H), 4.54 – 4.47 (m, 3H), 3.88 (s, 3H), 3.87 (s, 3H), 3.79 (t, J = 5.0 Hz, 2H), 3.64 – 3.51 (m, 12H), 3.43 (t, J = 5.9 Hz, 2H), 3.34 (t, J = 5.1 Hz, 2H), 3.29 – 3.21 (m, 4H), 2.66 (d, J = 4.6 Hz, 3H), 2.59 (s, 3H), 2.40 (s, 3H), 1.61 (s, 3H).

(S)-1-(5-chloro-4-((8-methoxy-1-methyl-3-(2-(methylamino)-2-oxoethoxy)-2-oxo-1,2-dihydroquinolin-6-yl)amino)pyrimidin-2-yl)-N-(3-(3-(2-(4-(4-chlorophenyl)-2,3,9-trimethyl-6H-thieno[3,2-f][1,2,4]triazolo[4,3-a][1,4]diazepin-6-yl)acetamido)propoxy)propyl)piperidine-4-carboxamide (**JWZ-7-97**)

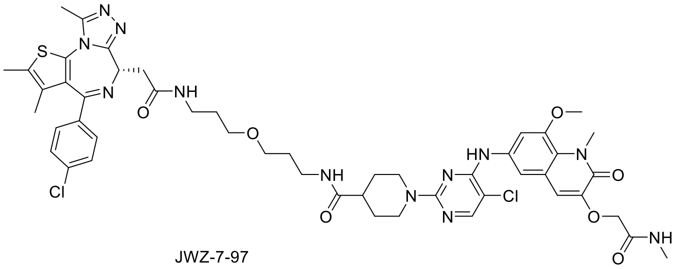

JWZ-7-97

**LC-MS** (ESI) m/z: 1027.4 $[M+H]^+$.

**1H NMR** (500 MHz, DMSO-d6) δ 9.10 (s, 1H), 8.15 (t, J = 5.7 Hz, 1H), 8.04 (s, 1H), 7.90 (q, J = 4.6 Hz, 1H), 7.75 (t, J = 5.7 Hz, 1H), 7.43 (d, J = 1.5 Hz, 2H), 7.41 (d, J = 8.8 Hz, 2H), 7.36 – 7.32 (m, 2H), 6.95 (s, 1H), 4.48 (s, 2H), 4.44 (dd, J = 8.1, 6.2 Hz, 1H), 4.36 (d, J = 13.1 Hz, 2H), 3.79 (s, 3H), 3.78 (s, 3H), 3.29 (dt, J = 24.4, 6.2 Hz, 4H), 3.22 – 2.98 (m, 6H), 2.84 (t, J = 12.9 Hz, 2H), 2.58 (d, J = 4.6 Hz, 3H), 2.52 (s, 3H), 2.32 (s, 3H), 1.67 – 1.56 (m, 4H), 1.53 (d, J = 2.6 Hz, 4H), 1.44 (qd, J = 12.5, 4.0 Hz, 2H), 1.16 (s, 1H).

(S)-2-((6-((5-chloro-2-(4-(2-(2-(2-(4-(4-chlorophenyl)-2,3,9-trimethyl-6H-thieno[3,2-f][1,2,4]triazolo[4,3-a][1,4]diazepin-6-yl)acetamido)ethyl)-2,7-diazaspiro[3.5]nonane-7-carbonyl)piperidin-1-yl)pyrimidin-4-yl)amino)-8-methoxy-1-methyl-2-oxo-1,2-dihydroquinolin-3-yl)oxy)-N-methylacetamide (**JWZ-7-98**)

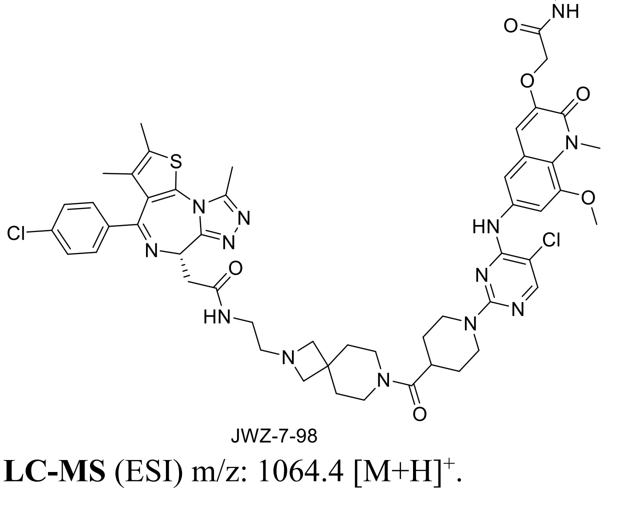

JWZ-7-98

**LC-MS** (ESI) m/z: 1064.4 $[M+H]^+$.

**1H NMR** (500 MHz, DMSO-d6) δ 9.07 (s, 1H), 8.14 (s, 1H), 8.04 (s, 1H), 7.90 (q, J = 4.6 Hz, 1H), 7.75 (t, J = 5.7 Hz, 1H), 7.43 (d, J = 1.5 Hz, 2H), 7.41 (d, J = 8.8 Hz, 2H), 7.36 – 7.32 (m, 2H), 6.95 (s, 1H), 4.64 – 4.62 (m, 1H), 4.50 – 4.44 (m, 1H), 4.40 (d, J = 13.3 Hz, 2H), 3.79 (dd, J = 8.4, 3.6 Hz, 2H), 3.71 (d, J = 6.6 Hz, 1H), 3.58 – 3.17 (m, 4H), 2.93 (q, J = 14.1 Hz, 1H), 2.57 (d, J = 4.6 Hz, 3H), 2.52 (d, J = 4.9 Hz, 3H), 2.35 (s, 3H), 1.78 (dd, J = 50.1, 24.2 Hz, 4H), 1.62 -1.52 (m, 6H), 1.44 (q, J = 12.6 Hz, 2H).

(S)-1-(5-chloro-4-((8-methoxy-1-methyl-3-(2-(methylamino)-2-oxoethoxy)-2-oxo-1,2-dihydroquinolin-6-yl)amino)pyrimidin-2-yl)-N-(3-((3-(2-(4-(4-chlorophenyl)-2,3,9-trimethyl-6H-thieno[3,2-f][1,2,4]triazolo[4,3-a][1,4]diazepin-6-yl)acetamido)propyl)(methyl)amino)propyl)piperidine-4-carboxamide (**JWZ-7-100**)

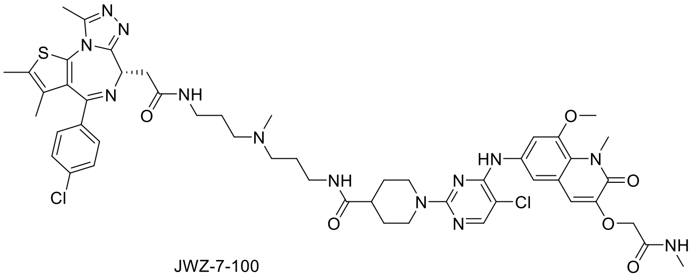

JWZ-7-100

**LC-MS** (ESI) m/z: 1040.4 $[M+H]^+$.

**1H NMR** (500 MHz, DMSO-d6) δ 9.47 – 9.33 (m, 1H), 8.94 (s, 1H), 8.32 (q, J = 6.2 Hz, 1H), 8.02 (s, 1H), 7.92 (dq, J = 18.7, 5.4, 4.8 Hz, 2H), 7.47 – 7.40 (m, 4H), 7.35 (d, J = 8.3 Hz, 1H), 6.94 (s, 1H), 4.48 (s, 2H), 4.46 – 4.39 (m, 2H), 3.80 (s, 3H), 3.78 (s, 3H), 3.28 – 3.11 (m, 4H), 3.09 – 2.80 (m, 10H), 2.67 (dd, J = 7.9, 4.8 Hz, 3H), 2.58 (d, J = 4.6 Hz, 3H), 2.53 (s, 3H), 2.32 (d, J = 3.6 Hz, 3H), 1.79 – 1.64 (m, 6H), 1.54 (s, 3H), 1.49 – 1.38 (m, 2H).

**Protein Constructs and Purification for TR-FRET**

The construct for 6xHis-BRD4_BD1 was described in Filippakopoulos, Qi et al[4] and was a gift from Nicola Burgess-Brown (Addgene plasmid # 38943 ; http://n2t.net/addgene:38943 ; RRID:Addgene_38943). Rosetta 2(DE3) (Sigma #71400) *E. coli* cells were transformed with plasmid and a single colony was inoculated as a starter culture in LB overnight. 25mL starter culture was added to 1L SB and grown to OD600=0.6 at 37 $^{O}$C, at which point 0.4mM IPTG (Sigma #I678) was added. Cells were harvested after 3.5hrs in 20mL lysis buffer containing 50mM Tris-HCl pH 8, 0.5M NaCl, 1mM TCEP, and 1mM PMSF by centrifuging at 6,000g for 15mins at 4 $^{O}$C. Lysate was incubated with 2mg chicken egg white lysozyme (Sigma #71412) and 500ug DNAseI (Sigma #DN25) for 30mins at 4 $^{O}$C, and lysed by high-pressure homogenization using an Emulsiflex (Avestin). After centrifugation twice at 10,000 rpm for 10mins at 4 $^{O}$C in a JLA25.50 rotor (Beckman), clarified lysate was incubated with 1mL bed volume TALON resin (Takara #635502) overnight at 4 $^{O}$C. Resin was washed with lysis buffer and a low-salt buffer + 10mM imidazole (50mM Tris-HCl pH 8.0, 25mM NaCl, 10mM imidazole, 1mM TCEP, 1mM PMSF) and eluted in the low-salt buffer containing 150mM imidazole. The elute was applied to a HiTrap HP Q 5mL anion exchange column (Cytiva #17115301) at 1mL/min and the flow-through collected. After concentration in a 3,000 MWCO Amicon filter (Millipore #UFC9003), the sample was applied to a Superdex 75 Increase 10/300 GL gel filtration column (Cytiva #29148721) equilibrated in storage buffer (10mM HEPES pH 7.4, 200mM NaCl, 1mM TCEP, 10% glycerol). Elutes were run on SDS-PAGE and those containing protein were stringently selected based on purity, pooled, concentrated to >250µM and flash-frozen in small aliquots at -80 $^{O}$C. A fresh aliquot was thawed for each assay.

The construct for BCL6_BTB-AviTag was based off previously designed BCL6 constructs used for TR-FRET assays, as reported in multiple papers including[5,6] and contains amnio acids 5-129 with three mutations, C8Q, C67R, C84N, that enhance stability but have no difference on backbone structure with the wild-type version[7]. The construct was ordered from Twist and was sub-cloned as a GST-thrombin site fusion protein in a pGEX vector and expressed similarly as the BRD4 construct. Clarified lysate was applied to 1mL bed volume of glutathione-superflow resin (Takara #635607) and incubated overnight at 4 $^{O}$C. After washing with lysis buffer containing 0.1% triton-X100 and no PMSF, the protein was cleaved off the resin by incubation for 3hrs at room temperature with thrombin (Sigma #T4648) in a buffer containing 20mM Tris-HCl pH 8.0, 150mM NaCl, 2.5mM $CaCl_2$, and 1mM TCEP. 5mM PMSF was added and the sample was immediately applied to a Superdex 75 Increase 10/300 GL gel filtration column equilibrated in 25mM Tris-HCl pH 8.5, 450mM NaCl, 1mM TCEP, and 10% glycerol. Fractions with protein were stringently selected for purity based on SDS-PAGE, pooled, concentrated to >150µM using a 3,000 MWCO Amicon filter (Millipore #UFC9003), and flash frozen and stored at -80 $^{O}$C. Biotinylation was carried out with 2.5µg homemade BirA (produced with pTP264, a kind gift from Dirk Görlich (Addgene plasmid # 149334 ; http://n2t.net/addgene:149334 ; RRID:Addgene_149334)) in a buffer with 40µM BCL6_BTB-Avi containing 50mM bicine pH 8.3, 0.05mM D-biotin (Sigma #2031), 10mM $MgCl_2$, and 10mM ATP for 1hr at 30 $^{O}$C. BirA was removed by incubation with His60 resin for 1hr at 4$^{O}$C. Biotinylation efficiency and protein purity was checked to be almost 100% by incubation of a small sample with streptavidin beads and monitoring of the flowthrough by SDS-PAGE.

**Protein Constructs and Purification for ITC and BLI**

The same procedure was followed for BRD4_BD1 as above except that after anion exchange, 1:100 TEV w/w was added to incubate overnight at $4^{O}C$, then desalted to remove the imidazole back into low-salt buffer, and the tag was cleaned up using Ni-NTA resin for 2hr rotating at $4^{O}C$ before applying to size exclusion chromatography.

For BCL6_BTB (C8Q C67R C84N), a construct was made without the AviTag using a pET48b(+) vector with Thioredoxin-6xHis-3C N-terminal tag. A similar procedure to the BRD4_BD1 purification was followed with the following modifications: lysis buffer: 50mM Tris pH 8/250mM NaCl/5% glycerol/0.5mM TCEP+1mM PMSF; protein was cleaved off the resin with addition of 3C protease overnight at 4C; flowthrough was collected and applied to a Sephadex S75 16/60 HiLoad in the same final storage buffer of 10mM HEPES pH 7.5, 200mM NaCl, 5% glycerol, 1mM TCEP. Elutes were run on SDS-PAGE and those containing protein were concentrated to 8-10mg/mL using 3K MWCO filter centrifuge and flash frozen. Protein concentrations were measured by Bradford.

### Supplemental Figure 1

a: Uncropped blots pertaining to Extended Figure 1b

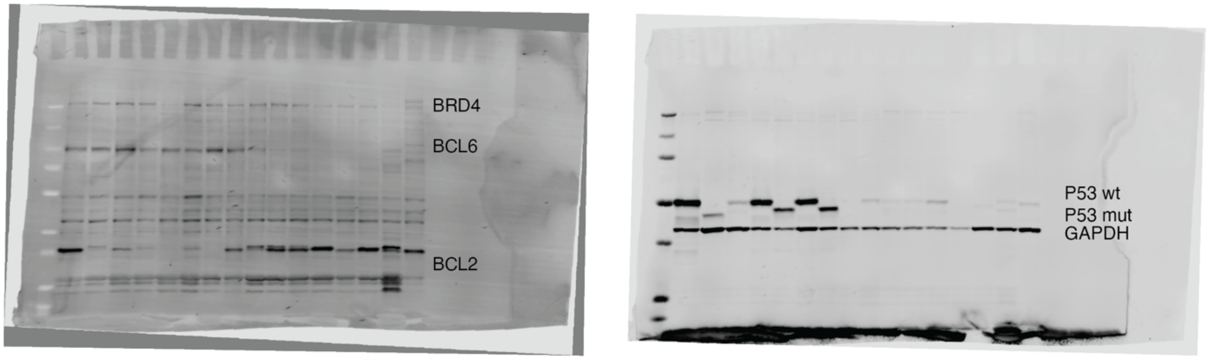


b: Uncropped blots pertaining to Figure 4d

KARPAS422

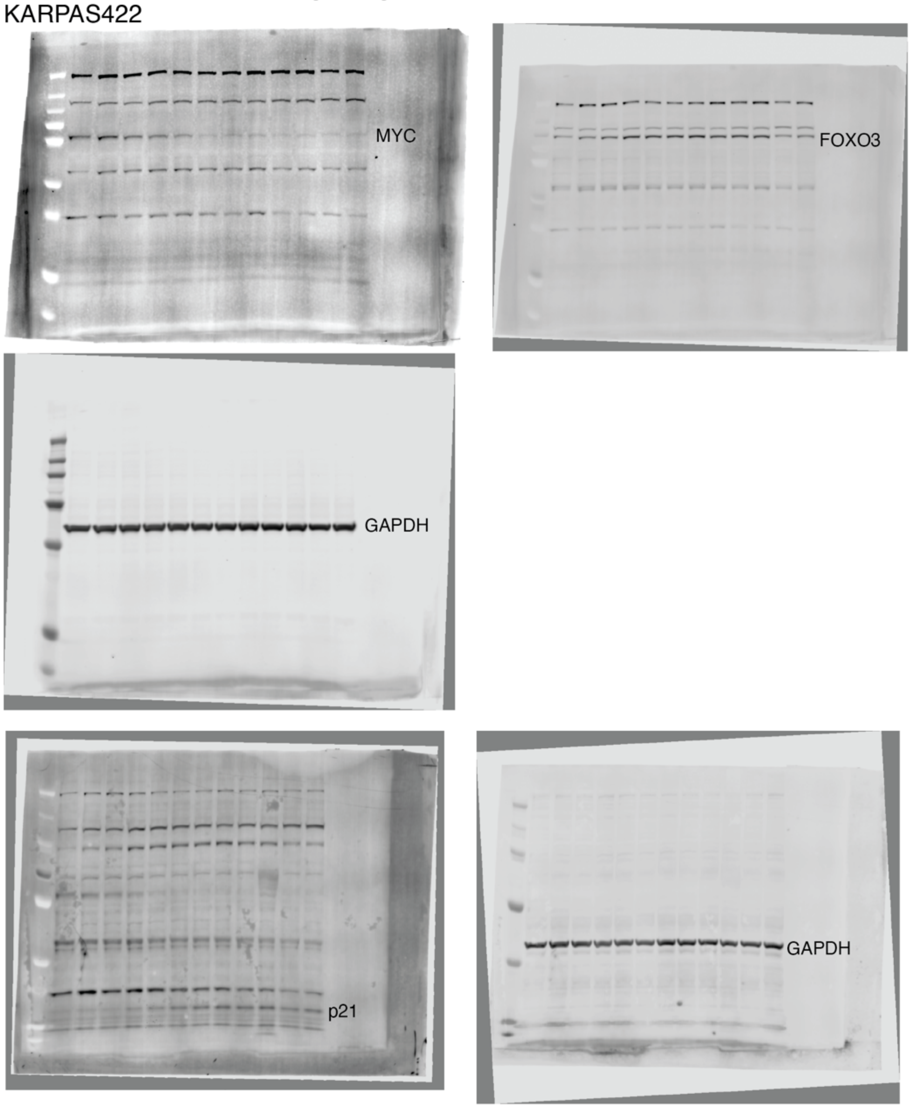

c: Uncropped blots pertaining to Figure 4d
SUDHL5

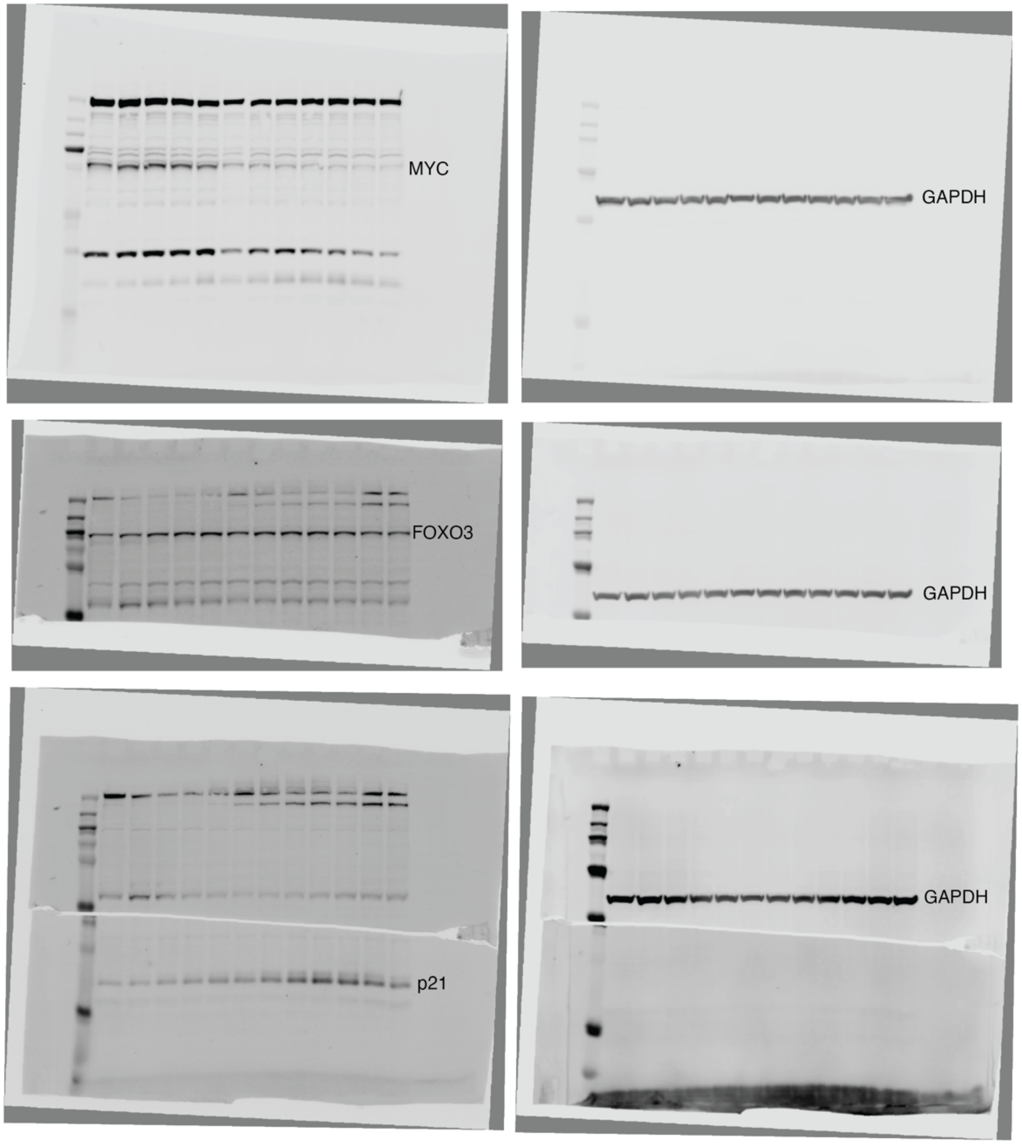

d: Uncropped blots pertaining to Figure 4e
KARPAS422

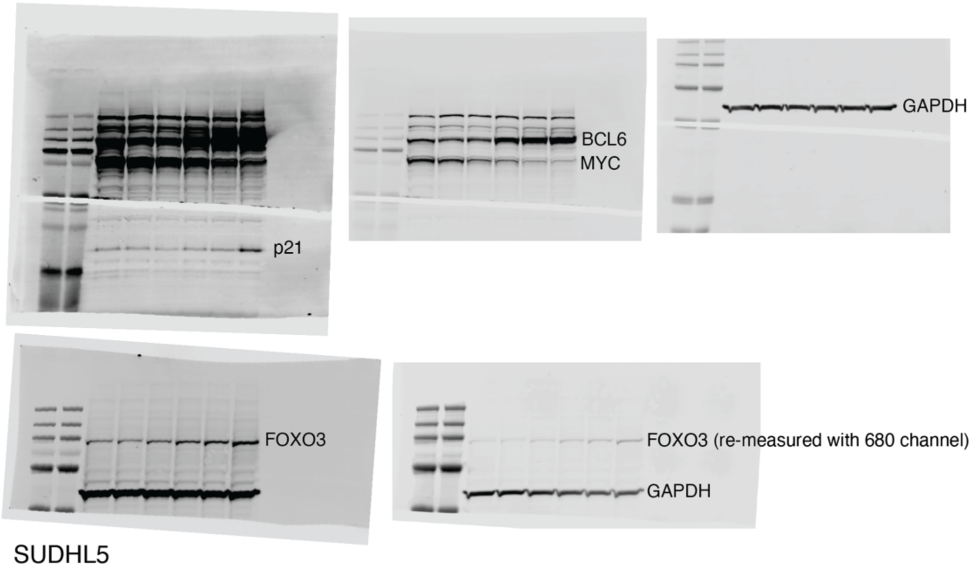


SUDHL5

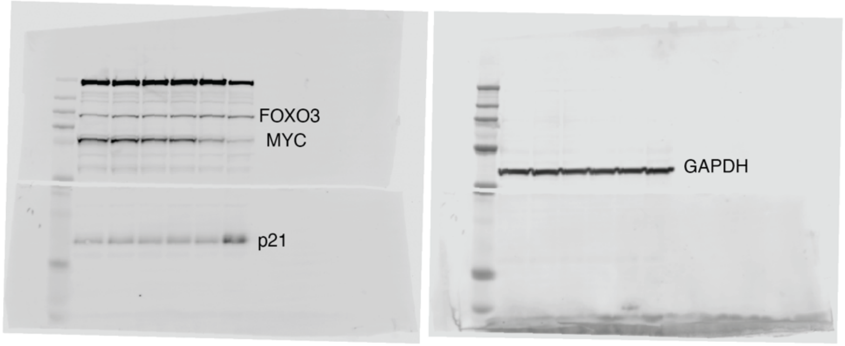


e: Uncropped blots pertaining to Figure 4g and Extended Data Figure 6b
KARPAS422

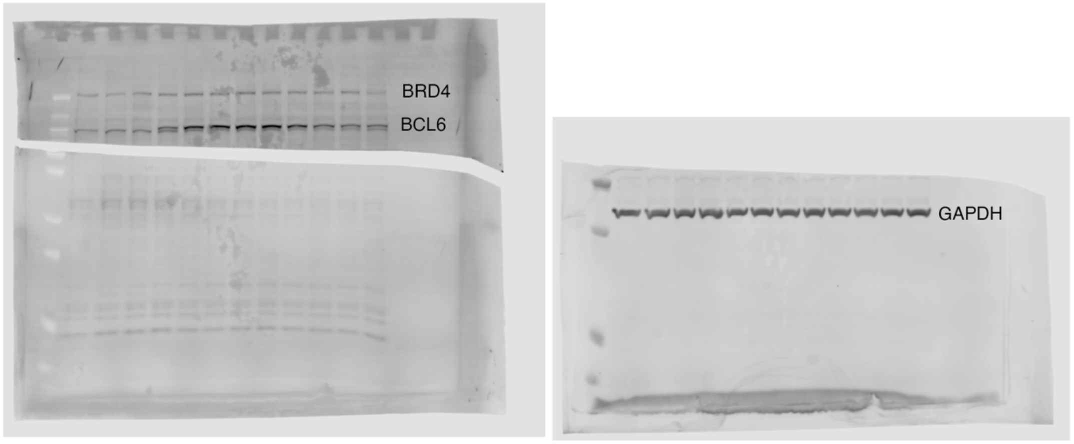

f: Uncropped blots pertaining to Figure 4g
SUDHL5

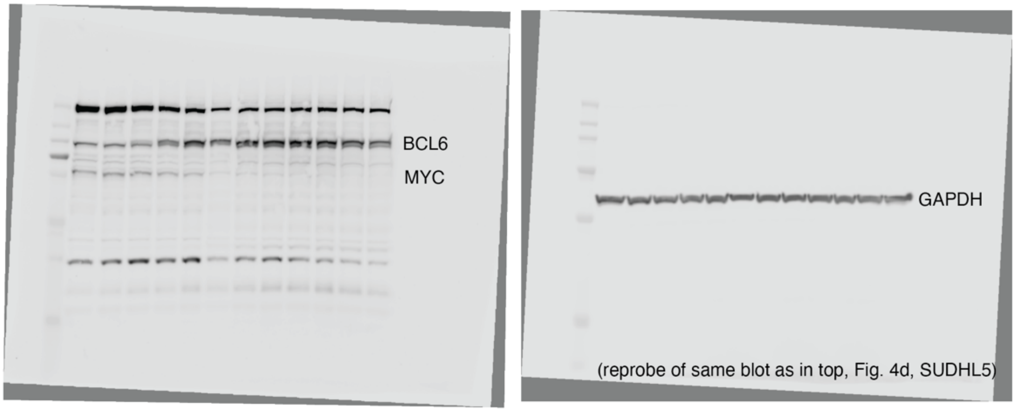


g: Uncropped blots pertaining to Figure 4i

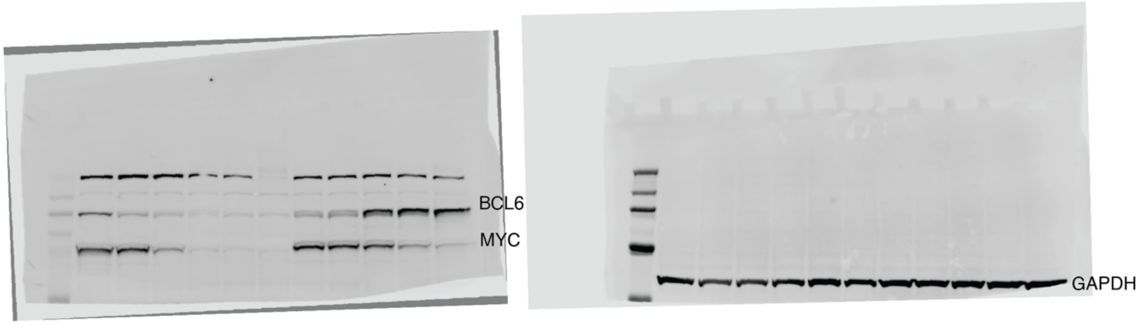


h: Uncropped blots pertaining to Extended Data Figure 5a
Neg1

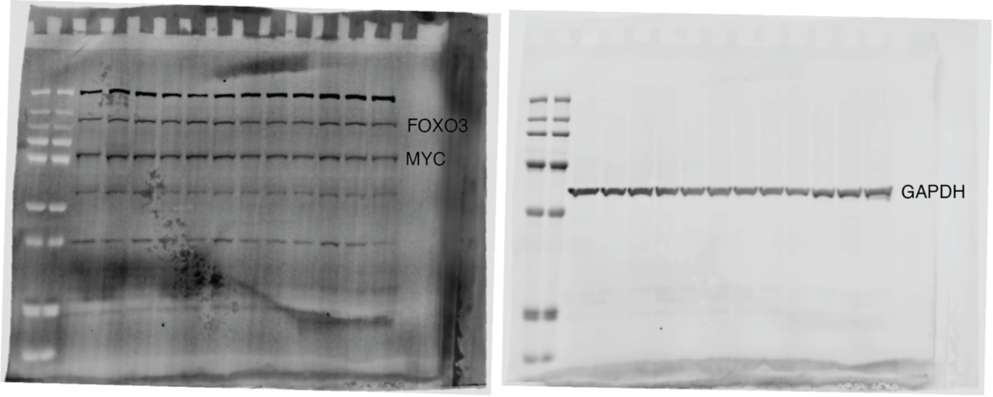

(cont.) h: Uncropped blots pertaining to Extended Data Figure 5a
Neg1

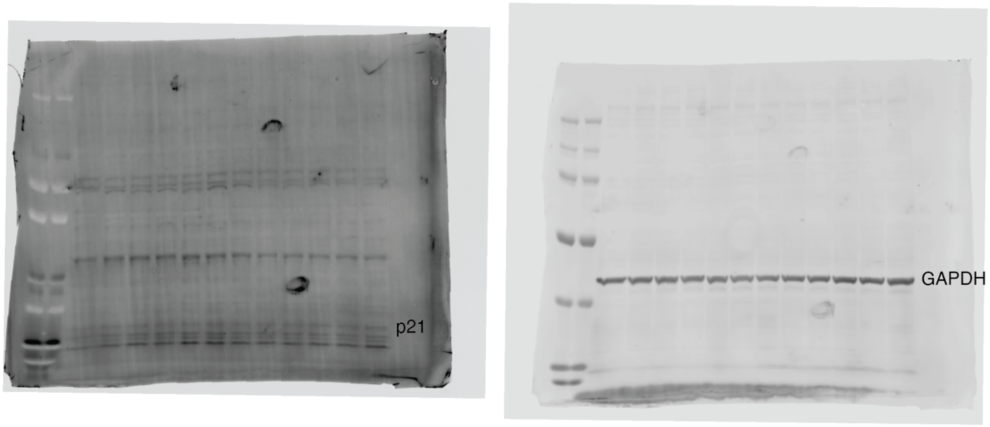


i: Uncropped blots pertaining to Extended Data Figure 5a
Neg2

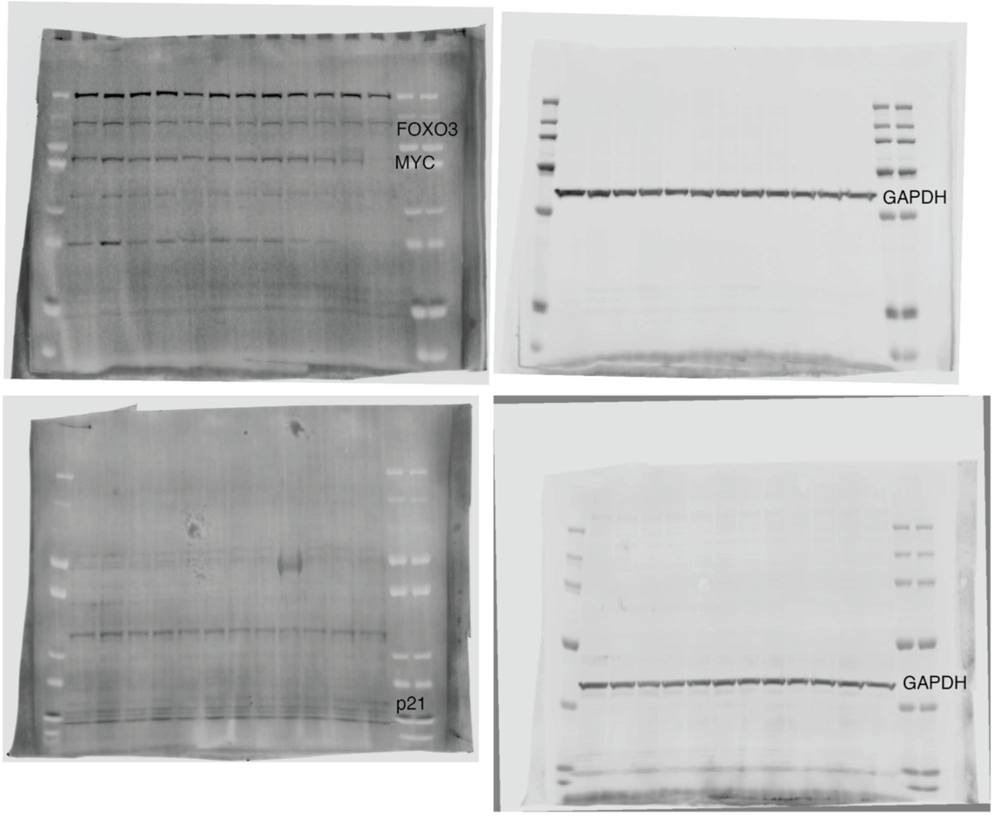

j: Uncropped blots pertaining to Extended Data Figure 6a
Neg1

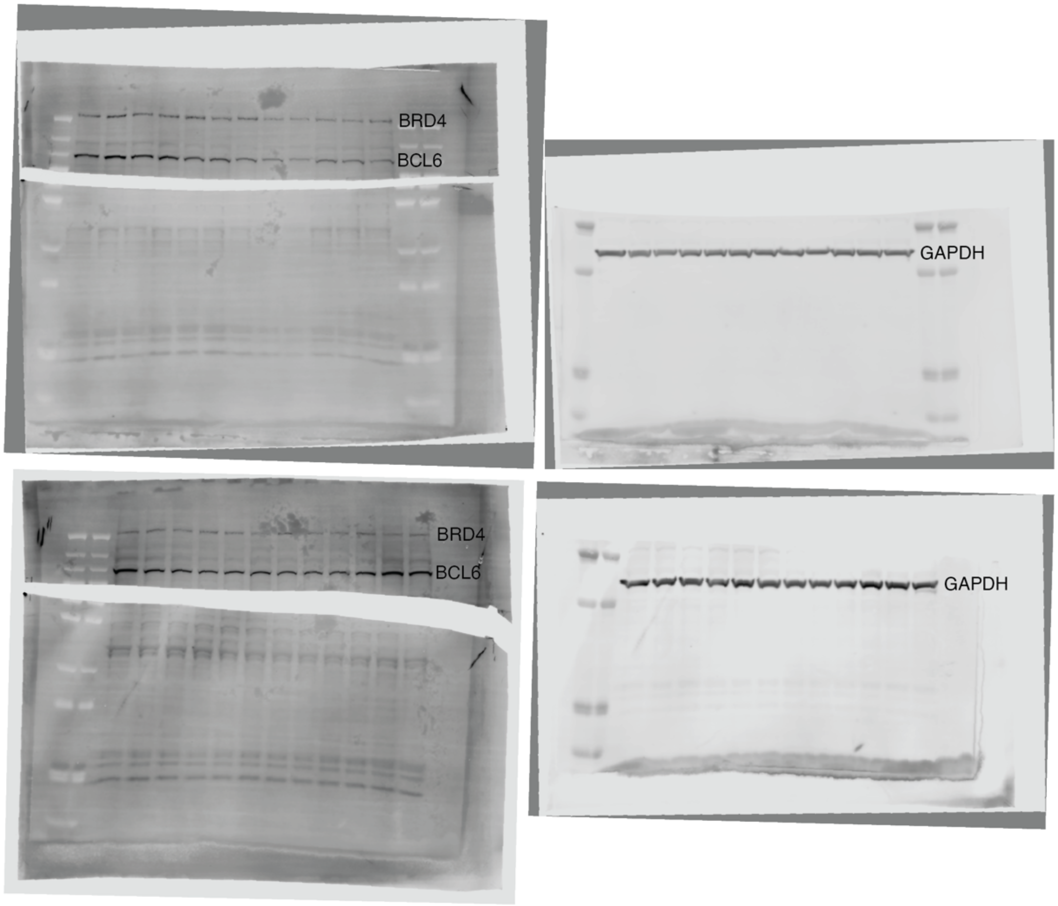


k: Uncropped blots pertaining to Extended Data Figure 6c
SUDHL5

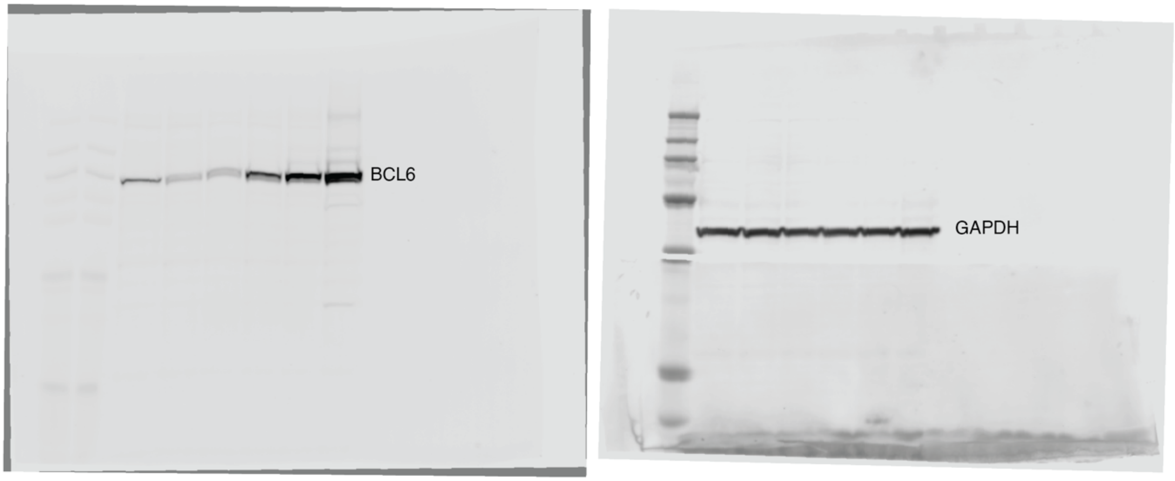

l: Uncropped blots pertaining to Extended Data Figure 6c
KARPAS422

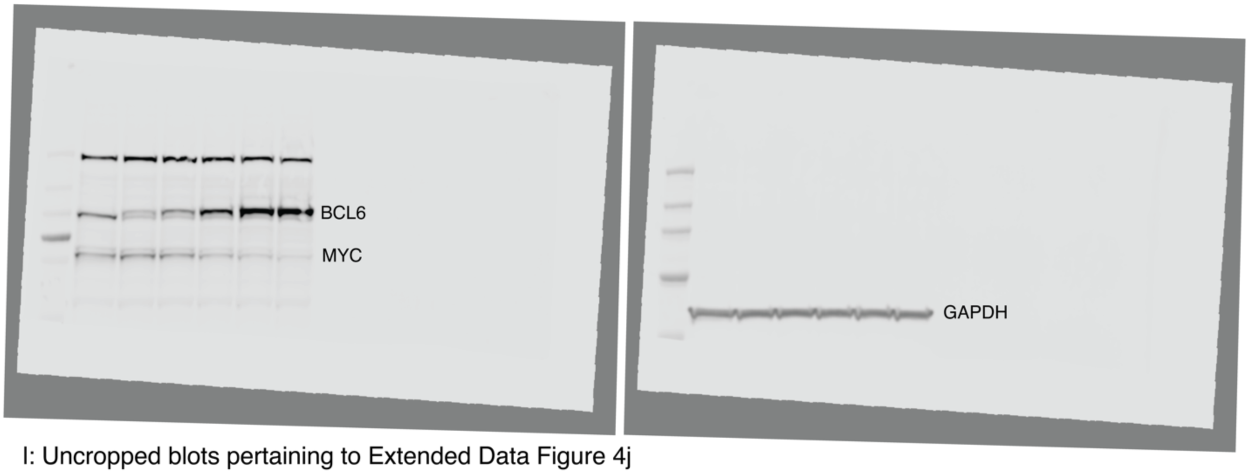


l: Uncropped blots pertaining to Extended Data Figure 4j
KARPAS422

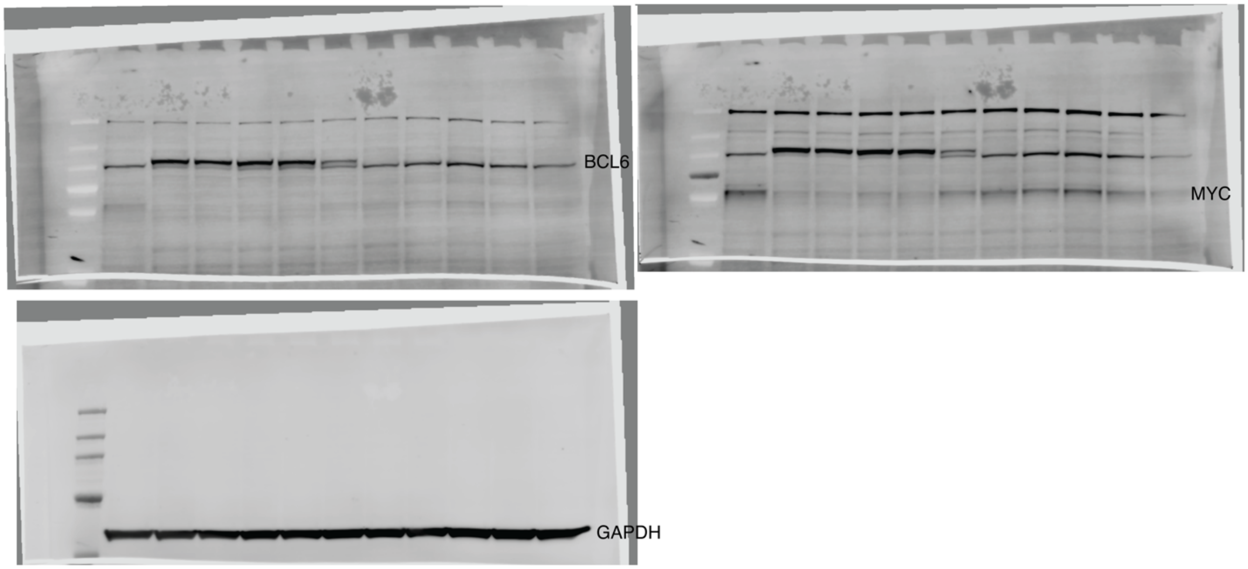

#### Supplemental Figure 2

a: Coomassie stained gel images of purified BRD4 and BCL6

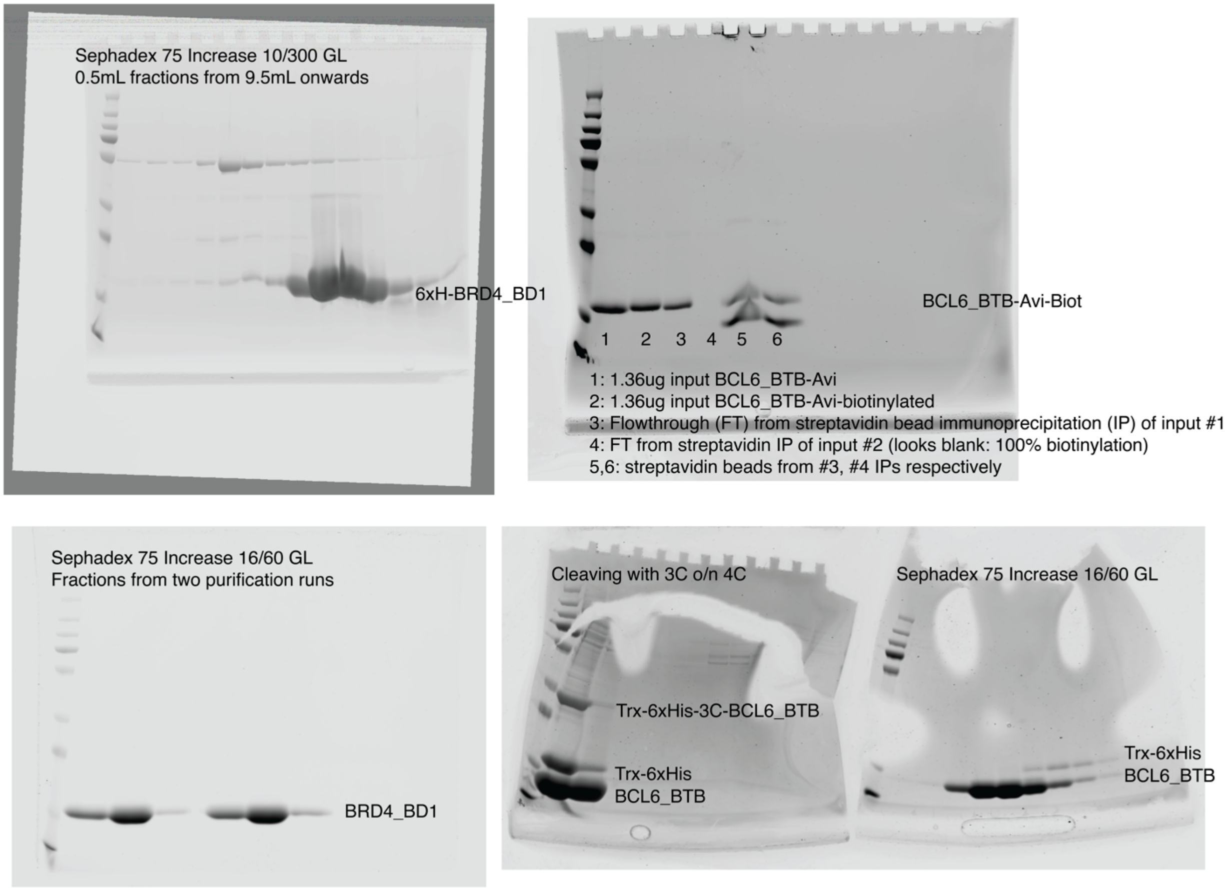

#### Supplemental Figure 3

a. Flow cytometry gating strategy for FITC-Annexin V/7AAD apoptosis assay

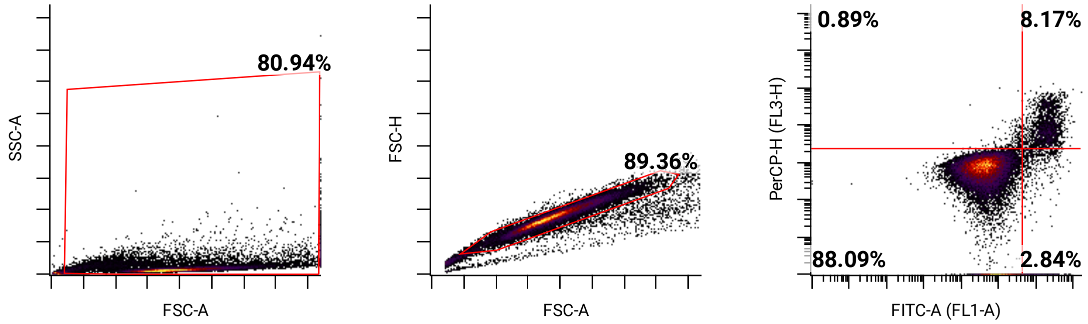


b. Flow cytometry gating strategy for cell cycle/TUNEL assay

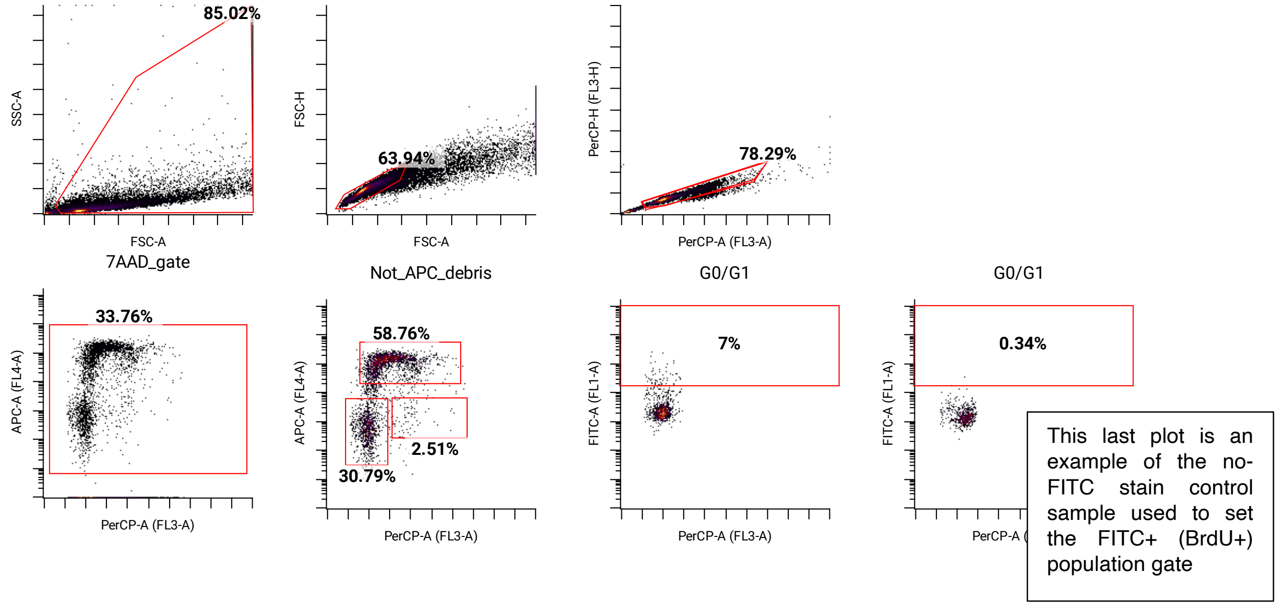
